# Robust probabilistic modeling for single-cell multimodal mosaic integration and imputation via scVAEIT

**DOI:** 10.1101/2022.07.25.501456

**Authors:** Jin-Hong Du, Zhanrui Cai, Kathryn Roeder

## Abstract

Recent advances in single-cell technologies enable joint profiling of multiple omics. These profiles can reveal the complex interplay of different regulatory layers in single cells; still, new challenges arise when integrating datasets with some features shared across experiments and others exclusive to a single source; combining information across these sources is called mosaic integration. The difficulties lie in imputing missing molecular layers to build a self-consistent atlas, finding a common latent space, and transferring learning to new data sources robustly. Existing mosaic integration approaches based on matrix factorization cannot efficiently adapt to nonlinear embeddings for the latent cell space and are not designed for accurate imputation of missing molecular layers. By contrast, we propose a probabilistic variational autoencoder model, scVAEIT, to integrate and impute multimodal datasets with mosaic measurements. A key advance is the use of a missing mask for learning the conditional distribution of unobserved modalities and features, which makes scVAEIT flexible to combine different panels of measurements from multimodal datasets accurately and in an end-to-end manner. Imputing the masked features serves as a supervised learning procedure while preventing overfitting by regularization. Focusing on gene expression, protein abundance, and chromatin accessibility, we validate that scVAEIT robustly imputes the missing modalities and features of cells biologically different from the training data. scVAEIT also adjusts for batch effects while maintaining the biological variation, which provides better latent representations for the integrated datasets. We demonstrate that scVAEIT significantly improves integration and imputation across unseen cell types, different technologies, and different tissues.

**Significance Statement:** Single-cell multimodal assays provide an unprecedented opportunity for investigating heterogeneity of cell types and novel associations with disease and development. Although analyses of such multimodal datasets have the potential to provide new insights that cannot be inferred with a single modality, access typically requires the integration of multiple data sources. We propose a probabilistic variational autoencoder model for mosaic integration, which involves merging data sources that include features shared across datasets and features exclusive to a single data source. Our model is designed to provide a lower dimensional representation of the cells for visualization, clustering, and other downstream tasks; accurate imputation of missing features and observations; and transfer learning for robustly imputing new datasets when only partial measurements are available.

## Introduction

With new technological advances, researchers are able to measure a growing number of molecular dimensions, including the genome, transcriptome and epigenome, on millions of cells. The primary goals are to classify subtypes of cells, to understand cell function, and to model basic biological processes such as early development and clinically relevant traits, such as disorders and cancer. Integrating data from multiple modalities [22] and whole-genome measurements presents new challenges in the analysis of single-cell data. While no single technology can measure all relevant omics in a single cell, recent developments facilitate the measure of several; for example, TEA-seq [29] and the DOGMA-seq [23] simultaneously measure chromatin accessibility, gene expressions, and protein abundances. However, obtaining single-cell multimodal datasets that measure many modalities may be costly and this limits the sample sizes. Moreover, there is a need to integrate new data sources with existing multimodal atlases and impute the missing biological modalities. Therefore it is of fundamental importance to develop methodologies that can perform integrative analysis and cross-modal translation on the full range of jointly profiled multimodal single-cell datasets.

To integrate single-cell multimodal datasets, most existing methods identify anchors either explicitly or implicitly. Depending on the choice of anchor, the integration methods can be divided into three categories: horizontal integration, vertical integration, and diagonal integration [2]. Horizontal integration methods, including Seurat v3’s CCA [28], Harmony [16], and LIGER [27], use common modalities and features as anchors to link datasets containing different cells. Vertical integration approaches combine different modalities from datasets measured across a common set of cells. Representative works include 1) Seurat v4’s Weighted Nearest Neighbor (WNN) [12], which identifies influential pairs of features based on the relative utility of each data modality and maps query datasets to the reference dataset based on shared variable features, and 2) totalVI [9], which models paired gene and protein measurements. On the other hand, there are methods that perform horizontal and vertical integration to combine different samples and modalities simultaneously. For example, MultiVI [3] models paired and unpaired measurements of gene and chromatin accessibility. Diagonal integration is considerably more challenging because neither modality nor cells are assumed to be shared. Extra cellular information is required to align different modalities.

As noted by Argelaguet et al. [2], a more general and challenging modern integration task is *mosaic integration*. For this integration task, different data modalities are profiled in different subsets of cells, or different subsets of cells are profiled from different experiments or technologies. Mosaic integration has two goals: (1) to map multimodal data to a common latent space, which is achieved by obtaining a joint multimodal profile for each cell, while utilizing the information from both shared and unshared features; and (2) to transfer knowledge of a fully trained model to a new dataset with partial modalities and features measured. This latter is sometimes also known as transfer learning [32, 34], and it is helpful in aligning cells and imputing unmeasured modalities and features for new datasets at a relatively low cost. Imputation of missing measurements and features is a natural bi-product of the integration procedure.

Recent advances in mosaic integration mainly focus on finding a common latent space while de-emphasizing the importance of imputation accuracy. For example, the non-negative matrix factorization algorithm UINMF [17] can align single-cell datasets containing both shared and unshared features in the low dimensional space; StabMap [10] first obtains low-dimensional score matrices for different datasets and then re-weights them based on features shared with the reference dataset. One common limitation of the current approaches is that only simple linear relationships between modalities are captured in the model [2]. And it is hard to incorporate different types of covariates for batch effect adjustment while retaining biological variation [31, 20, 7]. Besides, it is difficult for matrix factorization algorithms to align new datasets without training on the new dataset again. In the age of million single-cell multimodal data, both joint representations and transfer learning are indispensable. Although deep generative models such as totalVI [9] and MultiVI [3] can address the aforementioned limitations, they suffer from model misspecification issues when some features are missing for some cells. Specifically, existing deep generative methods either ignore unshared features or simply set the missing quantities to zero when performing integration, which induces biases. Failing to consider missing patterns, their performances will likely deteriorate when the proportion of unshared features increases significantly. As a result, new computational approaches that systematically model and utilize the information of missingness for combining mosaic-type multimodal datasets in the two scenarios are desirable.

We develop **S**ingle-**C**ell **V**ariational **A**uto**E**ncoder for **I**ntegration and **T**ransfer learning (scVAEIT), a probabilistic deep learning algorithm [15, 9, 8] capable of performing mosaic integration and imputation. The model allows for arbitrary patterns of shared and unshared features and modalities, and the integration does not require the input of any extra biological information. In addition to great flexibility, a primary advantage of the approach is its robustness to overfitting. By incorporating a masking procedure, our model learns interpretable joint representations for cells and the distributions of unobserved features conditioned on an arbitrary subset of observed modalities and features. Unlike the traditional generative models that are fully unsupervised, imputing the masked features serves as a supervised learning procedure, which is analogous to the supervised PCA in Seurat’s WNN [12]. The masking procedure also acts as a form of regularization to mitigate overfitting, which is typical of the deep generative models. In contrast, conventional neural networks only focus on and remember, for example, genes and proteins that are easy to predict in multitask learning. scVAEIT is thus extremely useful for missing features imputation and crossmodal generation, providing great flexibility and high accuracy in learning a common latent space. Furthermore, it can robustly transfer crossmodal knowledge to new single-modal and multimodal datasets that only measure partial panels of features of the training datasets, providing generalization on transfer learning.

## Results

### Method overview

An overview of the multimodal single-cell data analysis pipeline with scVAEIT is shown in Fig. 1. Existing single-cell data studies provide researchers with a variety of multimodal and single-modal datasets, illustrated here with three datasets of PBMCs (peripheral blood mononuclear cells) [23]: DOGMA-seq (RNA, protein, peaks), CITE-seq (RNA, protein), and ASAP-seq (RNA, peaks). Building on deep generative models, scVAEIT provides a flexible way to jointly analyze multiple multimodal and single-modal single-cell datasets while incorporating additional covariates for batch effect adjustments. After the model is trained on the mosaic-type dataset, it can then be applied to various downstream tasks. Specifically, it enables joint latent representations (intermediate integration) of all modalities and transfer learning to new data sources (late integration). Most importantly, it can robustly impute the missing quantities of the mosaic-type datasets. Though the model we illustrate in this setting is applicable for general multi-omic settings.

**Figure 1:**
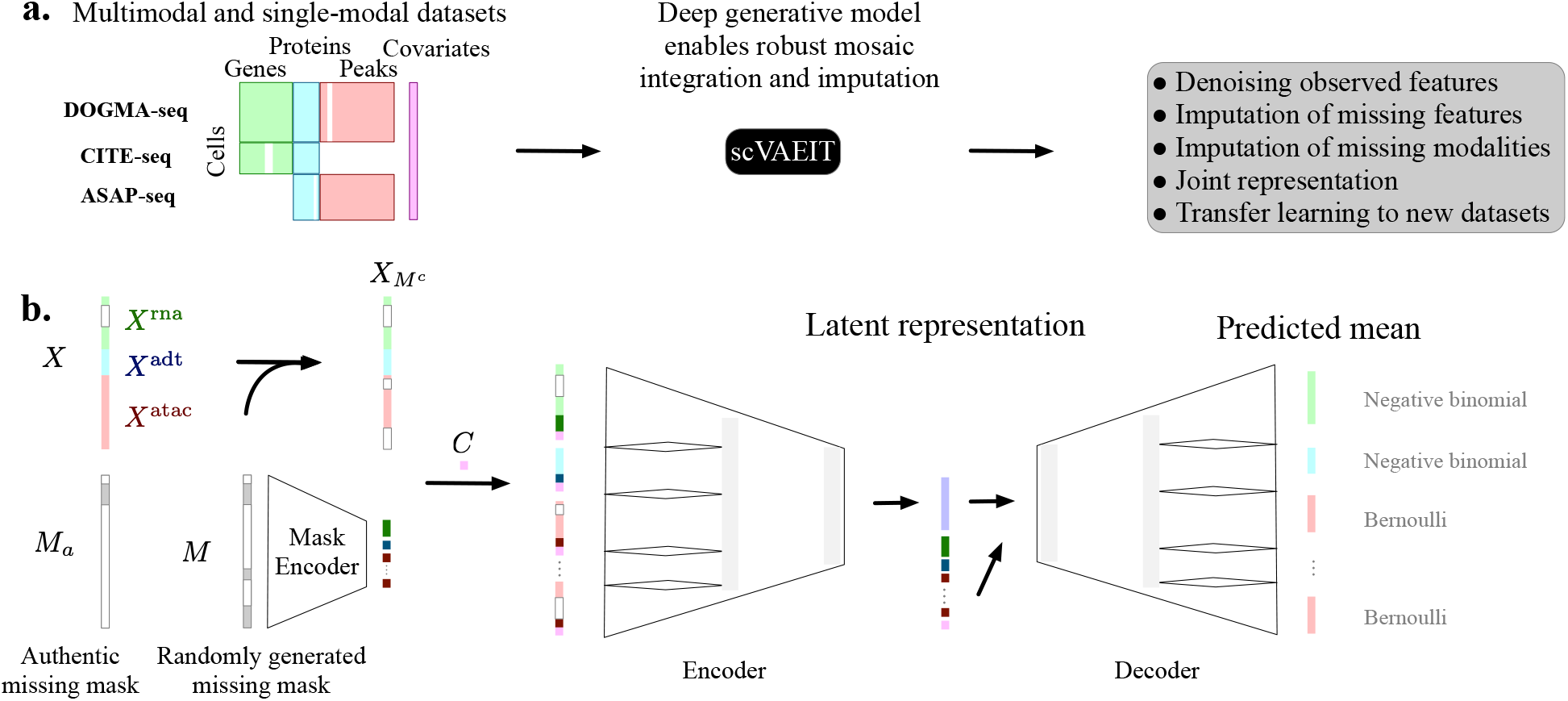
Overview of multimodal single-cell datasets mosaic integration with scVAEIT. **(a)** Multimodal sequencing techniques such as DOGMA-seq, CITE-seq, and ASAP-seq simultaneously measure multiple modalities in single cells. Different datasets can measure different panels of features for the same modality, producing mosaic count matrices for RNA, proteins and peaks. scVAEIT takes these matrices and an optional matrix containing continuous and categorical covariates to learn cross-modality and cross-feature relationships. It also produces a joint representation of all modalities for each cell. **(b)** For each cell, scVAEIT uses a mask to inform the missingness of the data. The actual or authentic mask is *M_a_*. During training, masks (*M*) are randomly generated and encoded for each modality to force the model to learn to predict specific portions of the features based on other observed features (*X_M^c^_*). In the first layer of the encoder and the last layer of the decoder, different modalities and peaks in different chromosomes break into subconnections that greatly reduces model complexity. The encoder then outputs the parameters of the posterior distributions for the latent variable *Z*, given the unmasked data *X_M^c^_* and the mask *M*. Next, samples from the posterior distributions along with the mask and covariates are fed to a decoder neural network to predict the posterior mean of *X*. Ultimately unobserved values are imputed and observed values are denoised.

The critical innovation of scVAEIT lies in the introduction of a mask *M*, incorporating information about missingness and the complementary observed features *X_M^c^_* (Fig. 1b). Instead of isolating shared and unshared features in multiple datasets, we utilize a missing mask *M* for each cell to inform scVAEIT about the missing pattern when performing mosaic integration. Hence, scVAEIT enables integrative analysis of cells from different sources with more or fewer features and modalities measured. It differs from other deep generative models in that it explicitly learns the conditional distributions of certain masked features (or modalities) given unmasked features (or modalities). In contrast, other variational inference models simply set the missing quantities as zero, which biases the learned models.

Even though we do not impose any assumption on the relationships between the shared and unshared features, scVAEIT learns the interdependence among features and modalities during the optimization process. This is achieved by sampling random mask *M* to force the model to predict the masked features based on the unmasked features. For example, masking out the peaks for cells in the DOGMA-seq datasets helps to impute the chromatin accessibility for cells in the CITE-seq datasets. This is also valuable for dealing with structural missing problems as we can exploit prior knowledge about data. On the other hand, randomly masking out a small portion of features also acts as a way of regularization, which prevents the model from overfitting the training dataset.

Because scVAEIT is optimized by the mini-batch stochastic gradient descent algorithm, its memory usage does not depend on the number of cells, which makes it scalable for large datasets. For instance, it can process the DOGMA-seq dataset with 13,763 cells and over 29,139 features (genes, proteins, and chromatin accessibility) within one hour for intermediate integration on a single Tesla v100-32 GPU. Once the model learns the cross-modality and cross-feature relationship from the training dataset, it can then robustly transfer its knowledge (late integration) to new sources at a relatively low cost. For example, denoising and imputing the previous DOGMA-seq dataset takes less than one minute on a single GPU. As more and more multimodal single-cell atlases become available, it is valuable to train scVAEIT on a reference atlas for once and readily transfer learns the new sources for cross-modality translation and imputation. Details of the specifications of the model architecture and training procedures are included in the “Method” section and Supplementary Note.

### Cross-domain translation with high accuracy

To examine cross-domain translation accuracy, we used a dataset consisting of PBMCs processed by CITE-seq [12] (see the “Methods” section). We held out one cell type for evaluation, trained different models based on the remaining cells, and then imputed each modality given the other for each held-out cell type. The two largest cell types – Mono (*n* = 49, 010 cells) and CD4 T (*n* = 41,001 cells) – were examined, and the results are summarized in Fig. 2a. As the held-out cell types were not present in the training set, high accuracy on cross-domain translation indicates that the model learns cross-modality relationships rather than memorizing the training set.

**Figure 2:**
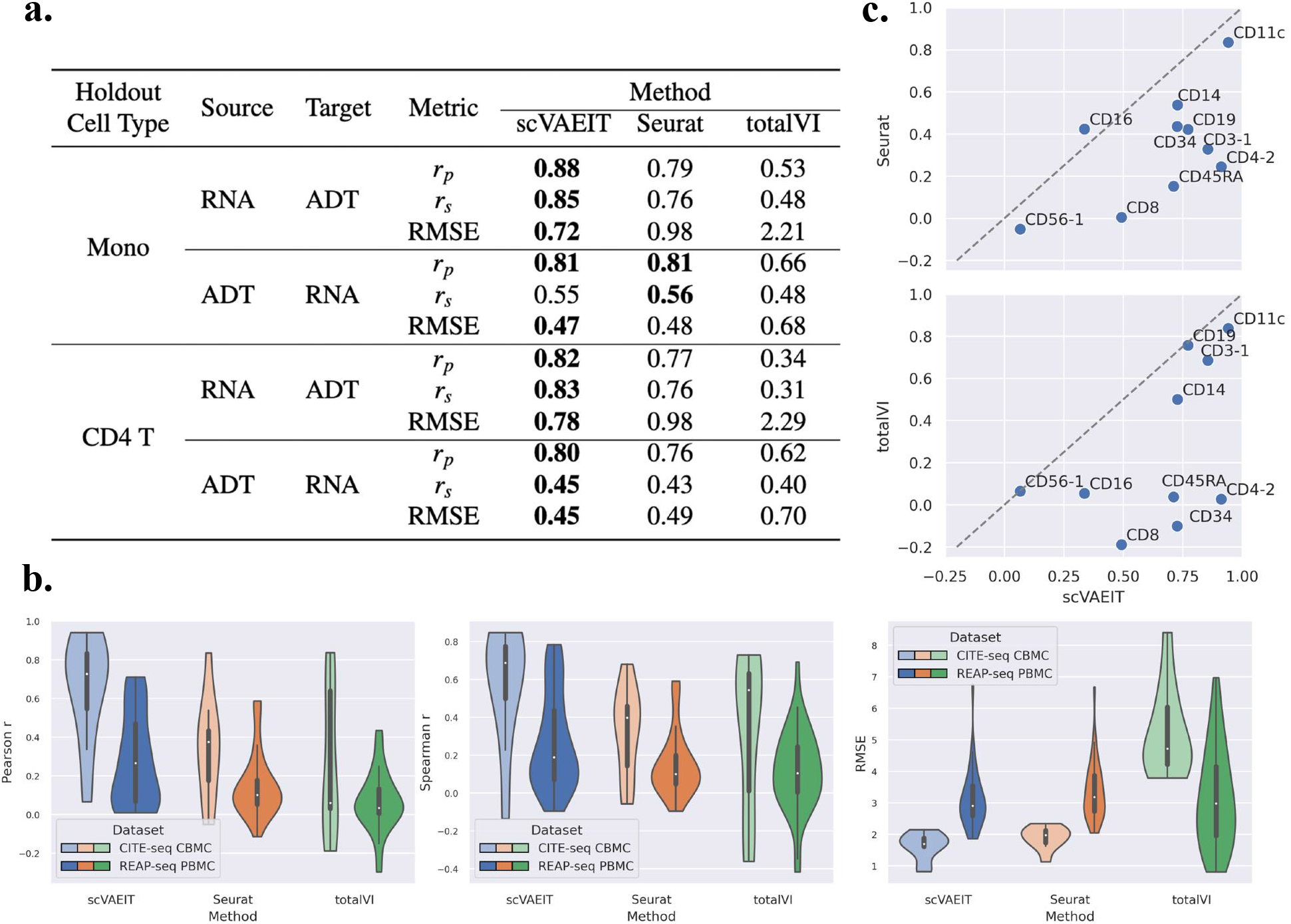
Evaluation of protein imputation on two external datasets. **(a)** Performance of missing modality imputation on the hold-out cell types when the source modality is fully observed. The evaluation metrics include Pearson correlation (*r_p_*), Spearman correlation (*r_s_*) and root mean square error (RMSE). RNA and ADT (antibody derived tag) represent the gene and protein modalities, respectively. **(b)** Violin plots of the Pearson correlation, Spearman correlation and root mean square error (RMSE) between the imputed and the true protein abundances on CITE-seq CBMC dataset and REAP-seq PBMC dataset. **(c)** Across tissue correlations between imputed and measured protein abundance on the CITE-seq CBMC data: scVAEIT versus Seurat and scVAEIT versus totalVI.

We compared the imputation accuracy with Seurat v4 WNN [12] with multimodal anchor transfer and totalVI [9], a deep generative model designed for CITE-seq datasets (see the “Methods” section). In almost all cases, scVAEIT achieved higher correlation and lower root mean square error (RMSE) compared to Seurat’s WNN method and totalVI. For protein imputation, Seurat v4 WNN failed to extrapolate well on the unseen cell type because it is based on protein-gene similarity in the training set instead of learning the nonlinear relationship between proteins and genes as scVAEIT. On the other hand, totalVI performed worst on predicting protein counts given gene counts, as it is designed for joint representation of proteins and genes and naturally fails to translate between the two modalities. Overall, the results indicate that scVAEIT captures the cross-modality relationships very well and is accurate for imputing missing modalities.

### Transfer learning external datasets

We next applied scVAEIT to impute proteins of two external datasets without any additional fine-tuning or training. Unlike the CITE-seq PBMC dataset we used to train the model, the CITE-seq CBMC (cord blood mononuclear cell) dataset was from a different tissue, while the REAP-seq PBMC dataset was generated from a different experimental protocol. Both of the two datasets measure genes and proteins as well. Such datasets are challenging because of the experimental biases and variances. Because both of the external datasets consist of a few proteins, it is unlikely any model can successfully impute gene expressions based on proteins. Therefore, we only inspected how well the models can infer the proteins in the external datasets when a partial panel of the genes is observed. On these external datasets, scVAEIT aligned more accurately with measured protein levels than Seurat and totalVI (Figs. 2c and S1). More specifically, scVAEIT achieved strong positive correlation (median Pearson correlation = 0.73, median Spearman correlation = 0.69) and small root mean square error of 1.70) on the CITE-seq CBMC dataset. At the same time, it was more stable than totalVI on imputation accuracy (Fig. 2b). We note that scVAEIT was not enforced to focus on only a few proteins in these external datasets, as it was trained on a panel of 227 proteins. The superior performance across tissues and technical protocols is due to the training scheme of random masking, which helps scVAEIT to give all-around attention to individual proteins.

### Trimodal integration and imputation

The recently proposed DOGMA-seq protocol revealed distinct changes in different modalities during native hematopoietic differentiation and peripheral blood mononuclear cell stimulation. However, most deep generative models are restricted to bimodal analysis. For example, totalVI [9] jointly models genes and proteins, and MultiVI [22] jointly models genes and chromatin accessibility. Although the existing method BABEL [33] uses separate encoders and decoders to model each modality and could be extended to trimodal analysis, it is still conceptually hard to align different modalities in the latent space. The difficulty of multimodal analysis lies in combining and balancing information from different modalities; thus, we would like to examine how scVAEIT performs in such cases.

To quantify how leveraging trimodal information improves dataset imputation, we compared our method with totalVI and MultiVI. A stimulation indicator was provided for all three methods as an extra covariate for batch effect correction. We also included 3-way weighted nearest neighbors (3-WNN) in Seurat v4 [12] as a benchmarking method, using Harmony [16] to integrate the stimulated and control data. We held out each cell type in the DOGMA-seq dataset as the test set and trained all models based on the remaining cells. Then the models were evaluated by imputing each modality given the other modalities (Figs. 3 and S2).

**Figure 3:**
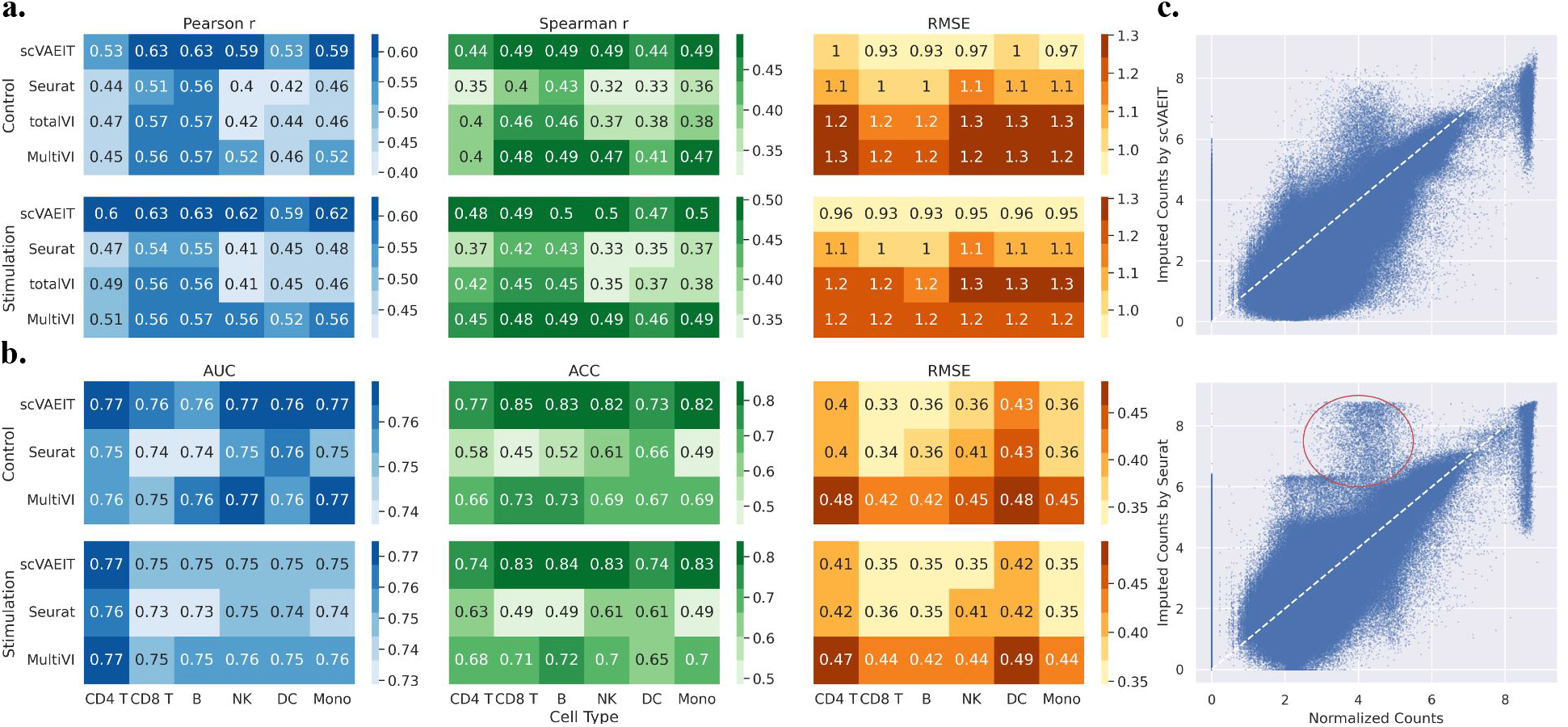
Trimodal imputation analysis on DOGMA-seq dataset. **(a)** Performance of gene imputation on the held-out cell types of the DOGMA-seq dataset. **(b)** Performance of chromatin accessibility imputation on the held-out cell types of the DOGMA-seq dataset. (The metrics AUROC and ACC denote the area under the receiver operating characteristic and accuracy, respectively). **(c)** Scatter plot of normalized protein counts versus imputed values on the held-out CD4 T cell type of the DOGMA-seq dataset by scVAEIT and Seurat. Each dot represents one protein abundance for one CD4 T cell. Because CD4 T cells in the query dataset are most similar to CD8 T cells in the reference dataset, Seurat simply memorizes the expression patterns of CD8a protein (within the red circle) from CD8 T cells and consequently overestimates its expression for CD4 T cells.

scVAEIT achieved strong performance across different held-out cell types and experimental conditions. For gene imputation, other deep generative models could not extrapolate well on the unseen cell types, producing large RMSEs, while Seurat’s anchor transfer method failed to capture the relationships between different modalities, resulting in low Pearson correlations and Spearman correlations (Fig. 3a). scVAEIT, however, excelled in both aspects for imputing the gene counts. The binary versus continuous nature of the three modalities required us to use different metrics for evaluating chromatin accessibility predictions. Inferring peaks from gene and protein expressions, it is on par with MultiVI on the area under the receiver operating characteristic (AUROC) while significantly better on accuracy (ACC) and RMSE (Fig. 3b). If we zoom in to look at the density scatterplot of the normalized protein counts and the imputed counts (Fig. 3c), we see that Seurat overestimated the protein abundances of CD8a in the held-out CD4 T cell type because the test cell type is similar to the CD8 T cell type, where the CD8a protein highly expresses. For imputation of sparse counts of genes MAML2, LINC00681, SOS1, FHIT, and MYBL1, we also observed Seurat’s underestimation and over smoothing (Fig. S3). These results indicate scenarios where Seurat’s map and query method might fail. On the contrary, scVAEIT learned a nonlinear and more accurate mapping from the expressions of all available modalities and features to the unknown features and hence achieved a more reasonable imputation on the missing quantities even in the unseen CD4 T cell type.

### Robustness to missing features

As more and more single-cell multimodal atlases become available, we could expect that late integration would be increasingly essential and practical for researchers to transfer knowledge from the atlases to the new datasets; however, the new datasets may not contain all the features measured in a reference atlas, leading to missing data issues in practice. Therefore, we investigated how different levels of missingness of the measured features might impact the late integration of new datasets by using the DOGMA-seq dataset. In the largest CD4 T cluster, a specific portion of genes, proteins, and chromatin accessibility, was randomly held out as a test set, and different models were trained on cells in other cell types and then applied to impute these missing quantities based on the observed features in the held-out cell type. The process was repeated multiple times.

Except for scVAEIT, we observed that other deep generative models have unstable behaviors on imputation accuracy under the model misspecified scenario (Figs. 4a-b for proteins and chromatin accessibility imputation, and Fig. S4 for genes imputation). More specifically, the performance of totalVI on gene and protein imputation deteriorated dramatically when the missing proportion increased; MultiVI, on the other hand, had a larger variance and uncertainty when facing a larger degree of missingness. On the other hand, as a nonparametric method, Seurat was more stable with respect to different levels of missingness (Fig. 4a). It obtained better Pearson correlations for gene imputation but worse Spearman correlation than MultiVI (Fig. S4), meaning that Seurat did not capture nonlinear relationships well among sparse signals. As noted, scVAEIT effectively combined the advantages of both methods, and was robust and accurate even when features were missing and the training model was misspecified to the new datasets.

**Figure 4:**
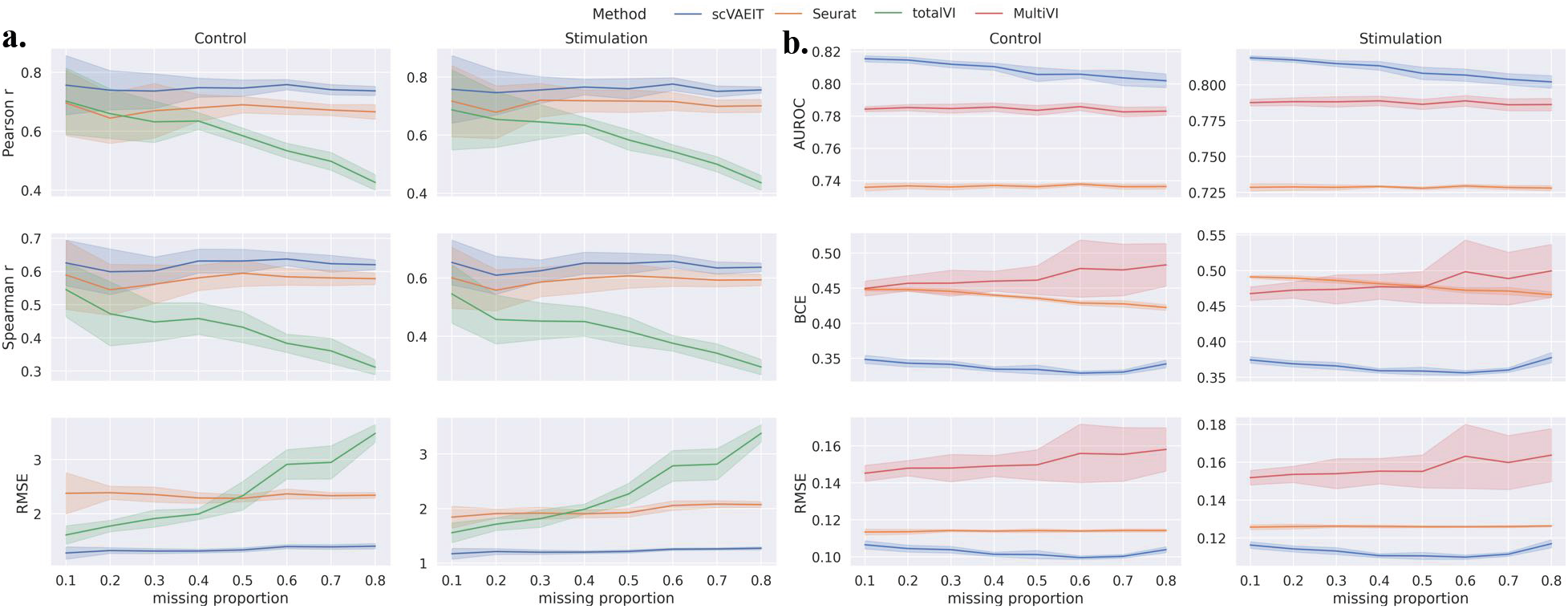
Incorporating masking information during training enables robust late integration with missing features. **(a)** Performance of protein imputation on the held-out CD4 T cell type of the DOGMA-seq dataset with random missing. For each missing proportion, all methods are evaluated using the same observed features across 10 runs and the shaded region is within one standard deviation of the average performance. **(b)** Performance of chromatin accessibility imputation on the held-out CD4 T cell type of the DOGMA-seq dataset with random missing (the metrics AUROC and BCE denote the area under the receiver operating characteristic and binary cross entropy, respectively). For each missing proportion, all methods are evaluated using the same observed features across 10 runs and the shaded region is within one standard deviation of the average performance.

### Application to multi-source and multimodal mosaic integration

We further applied scVAEIT to integrate the DOGMA-seq dataset with a CITE-seq PBMC dataset and an ASAP-seq dataset from Mimitou et al [23]. After filtering low-quality cells and features (see the “Method” section), the three datasets have 208 proteins in common, while the DOGMA-seq and the CITE-seq datasets have only 880 shared genes, and the DOGMA-seq and the ASAP-seq datasets have only 26,206 shared peaks (Table S1). We first considered the task of two-phase mosaic integration, where the mosaic multimodal datasets were combined through intermediate integration to remove the effects of experiment conditions, and the new mosaic multimodal datasets were imputed afterwards (late integration). Each cell type from the three datasets was held out for imputation and evaluation in turns, while the rest were used to perform intermediate integration. As all datasets measure a shared panel of proteins, it is easier to use protein counts to link these datasets together. Instead, we inspected how protein counts can be imputed based on other modalities in the held-out cell type.

We compared scVAEIT with Seurat’s 3-WNN and anchor transfer method. Although the procedure of two-phase integration is straightforward for scVAEIT, it becomes much more complicated for Seurat’s method. First, the multimodal reference and query datasets need to be integrated with Harmony to adjust for the stimulation effects separately, when performing dimension reduction. Then the query dataset is mapped to the reference dataset in the low-dimensional space. Finally, the missing quantities are imputed based on the similarity between the reference and query cells computed in the low-dimensional space. In our experiments, the multimodal neighbor method failed for integrating cells in the DC or Mono cell type because there were too few cells, no matter how we chose the number of neighbors. As shown in Fig. 5a, scVAEIT achieved consistently better performance on different cell types in terms of Pearson correlation, Spearman correlation, and RMSE. The overall better performance across the three different multimodal datasets also indicated that scVAEIT learns to infer protein counts based on gene expressions (CITE-seq), chromatin accessibility (ASAP-seq), or both (DOGMA-seq).

**Figure 5:**
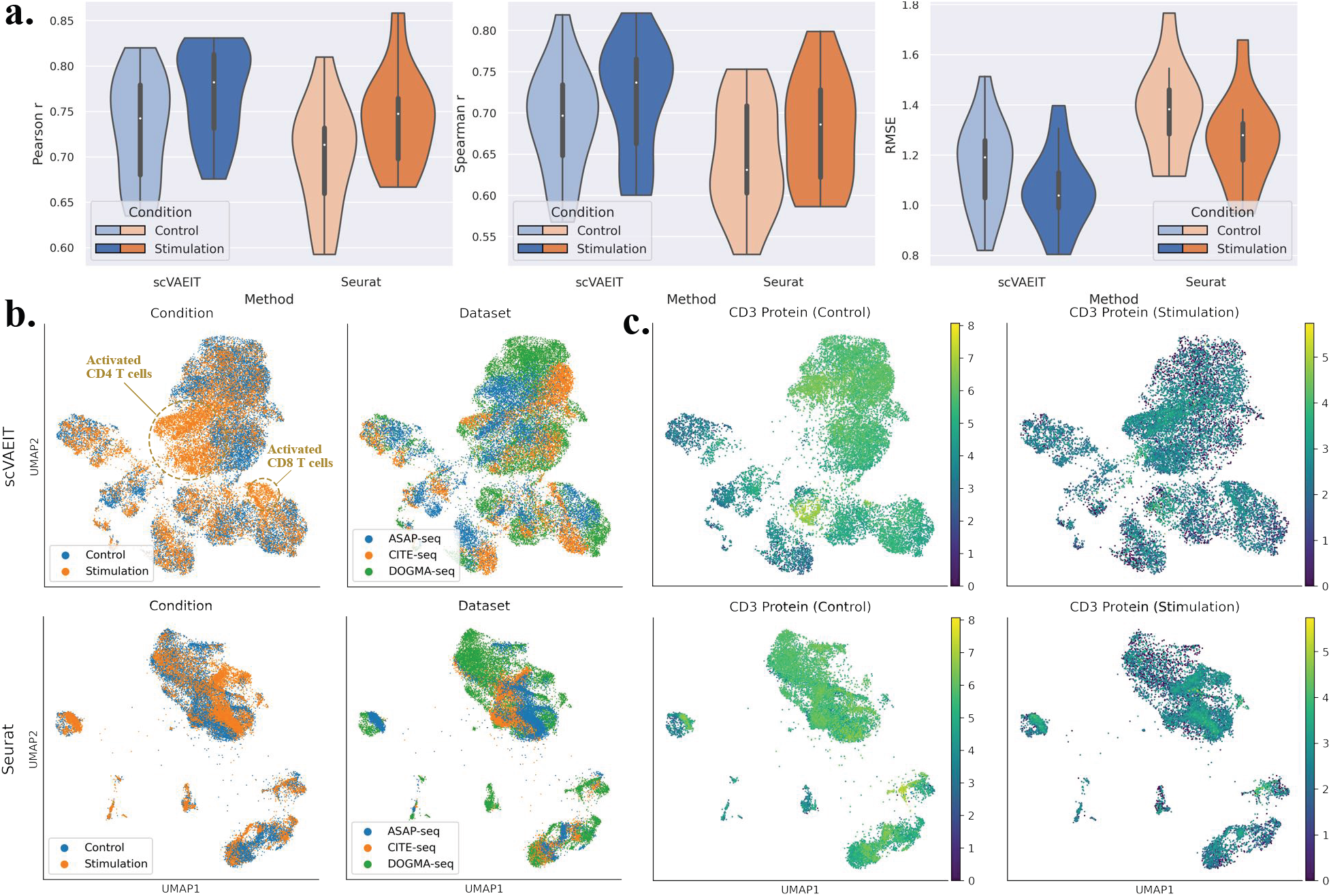
Integration of DOGMA-seq, CITE-seq, and ASAP-seq PBMC datasets. **(a)** Performance of protein imputation after two-phase integration with held-out cell types of three multimodal datasets. The two smallest cell types of the three datasets – DC and Mono – contain too few cells for Seurat’s integration method to find multimodal nearest neighbors, even with a reduced number of multimodal neighbors. **(b)** Joint embeddings of the three multimodal datasets after intermediate integration. Seurat’s WNN with Harmony overcorrects the experimental condition effect, eliminating the difference between the stimulated and control T cells. **(c)** Log-normalized expressions of CD3 protein in the control cells (first column) and the stimulated cells (second column) on scVAEIT’s and Seurat’s embeddings. Purple and yellow dots correspond to lowly and highly expressed cells, respectively.

Next, we refitted both models using all cell types and visualized the cell embeddings in Fig. 5b. We also compared with matrix the factorization method UINMF [17] (Fig. S5a). We performed Uniform Manifold Approximation and Projection for Dimension Reduction (UMAP) [21] directly on the learned latent variables for scVAEIT, while a similar UMAP visualization based on the trimodal WNN graph was also shown for Seurat. Fig. S6a shows the same embeddings colored by annotated cell types obtained from the original paper [23]. The majority of the CD4 T and the CD8 T cells, as identified in the original paper, reside in the upper right half and the lower right half of each cluster in Fig. 5b, respectively. Because a portion of the cells in all datasets was stimulated with anti-CD3/CD28, the stimulation-dependent changes within T cells can be observed. More specifically, the activated CD4 T and CD8 T cells clusters (clusters 5 and 16 in Fig. 5g of the original paper [23]) can be easily recognized in scVAEIT’s embeddings. On the other hand, integration based on Harmony and Seurat’s WNN removed not only specific batch effects but also meaningful biological signals: notably, it is not easy to identify stimulated T cells from Seurat’s embeddings; UINMF failed to adjust for batch effects correctly. By contrast, scVAEIT retained meaningful biological differences when adjusting for the effect of experimental conditions. When we visualized the source of datasets, we observed that scVAEIT merges cells from different datasets more evenly than Seurat. In scVAEIT’s embeddings, the cells from different datasets were aligned based on the expression of features; hence, they were not entirely mixed because of the differences in expression distribution from different techniques. If the researchers assure that such an effect should be eliminated, one can further apply batch effect correction methods such as FastMNN [11] on the latent variables and map all cells to the same source.

The dynamic changes in CD3, CD279, and CD69 protein abundances of control and stimulated cells were also examined in Fig. 5c, Fig. S5b and Fig. S6b-c. CD3 protein is highly expressed (green) in the CD4 T cell type (the top right cluster) and the CD8 T cell type (the bottom right cluster) in the absence of stimulation, while the differences between cell types become smaller in the stimulated cells, as expected. In the major CD4 T cell type, we noticed that the expression level of control cells gradually varied in scVAEIT’s embeddings while it remained almost the same in Seurat’s embeddings. Similar results can also be observed for CD279 and CD69 proteins (Fig. S6b-c), which have different expression patterns in the control and the stimulated cells. Furthermore, scVAEIT also provided a convenient way to visualize both denoised and imputed gene expressions, especially when a gene is only measured in some of the datasets (Fig. S7). For example, the CD3E gene is only available in the CITE-seq dataset (Fig. S7a), while the CD69 gene is only available in the DOGMA-seq and CITE-seq datasets (Fig. S7b). The denoised and imputed expressions can then be used to test and identify differential features on the integrated dataset (Fig. S10). Furthermore, this procedure can be naturally generalized for differential expression testing between case and control groups by applying the Bayesian inference framework [18, 5].

## Discussion

scVAEIT is a highly adaptable procedure, designed for multiple integration tasks, including joint representations (intermediate integration [22]), and transfer learning and imputation on new datasets (late integration [22]). First, scVAEIT utilizes information from shared and unshared features of different modalities, such as genes, proteins, and chromatin accessibility, to build an integrative probabilistic model for intermediate integration. The proposed model learns complex nonlinear relationships between modalities, enables a common latent representation of different modalities for each cell, and can accurately impute the missing quantities. Second, after training on some data source, scVAEIT readily transfers its knowledge to a new data source, even when the new dataset only contains partial measurements of features or modalities of the training data, enabling robust cross-modality translation. Third, scVAEIT is flexible in incorporating different types of covariates and adjusting for batch effects when combining single-cell datasets from various experiments, tissues, and technologies. Surprisingly, even including single-modal datasets can help with multimodal learning for scVAEIT, as revealed in Fig. S9. Compared to other deep generative models, the success of scVAEIT relies on the masking procedure that helps the model predict a certain portion of features in a supervised manner. This not only makes great usage of mosaic-type datasets but also serves a role of regularization.

The success of deep generative models [9, 3, 33] on single-cell multimodal data analysis enables expressive and scalable probabilistic representations of cells. Such models can harness the strengths of each modality by combining multiple cellular views in an end-to-end pipeline. The deep neural networks are adequate to describe complex and heterogeneous relationships between cells and modalities, and the probabilistic modeling provides interpretability with uncertainty quantification. Despite these promising results, the deep generative models can suffer from model misspecification issues when facing mosaic datasets. This happens when a model is used to perform intermediate integration on multiple multimodal datasets with different missing measurements or during late integration when a learned model is applied to transfer knowledge to new datasets that measure different panels of features from the training dataset. In experimental results, when the degree of model misspecification increases, the accuracy and stability of other deep generative models can be hugely impacted, and their performance can be inferior to nearest neighbor methods if the information of missingness is not taken into consideration.

During the training process of scVAEIT, the masking procedure plays a vital role in learning conditional distributions under arbitrary missing patterns. The supervised nature and regularization effect of the masking procedure produce more robust model constructions when learning from mosaic datasets and transferring knowledge to new datasets. There is still flexibility in choosing masks for training. For example, we can incorporate specific structural missing patterns to generate the masks with prior information on the predictable relationship between features. It will be valuable when we try to efficiently and systematically integrate more and more modalities.

## Methods

### Datasets preprocessing

Table S1 summarizes the information of the preprocessed datasets. For the CITE-seq PBMC dataset[12], we first identified the top 5000 variable genes and then filtered out genes and proteins expressed in less than 500 cells. The filtered dataset consists of 161,764 cells with 4,686 genes and 227 proteins. For the CITE-seq CBMC dataset[26], we first filtered cells that have less than 200 genes and then filtered out genes that are expressed in less than 10 cells. The filtered dataset consists of 7891 cells with 3464 genes and 10 proteins that are shared in the CITE-seq PBMC dataset. For the REAP-seq PBMC dataset[24], we first filtered cells with less than 200 genes or more than or equal to 5% mitochondrial counts and then filtered out genes expressed in less than 500 cells. The filtered dataset consists of 7,092 cells with 3,864 genes and 38 proteins shared in the CITE-seq PBMC dataset. For genes and proteins in the DOGMA-seq PBMC dataset, we applied the same procedure as above, resulting in 2,166 genes and 208 proteins. For chromatin accessibility, we retained the 75% variable peaks and filtered out peaks that appear in less than 500 cells or are on sex chromosomes, leaving 26,765 peaks. The CITE-seq PBMC and the ASAP-seq PBMC datasets from Mimitou et al.[23] are filtered analogously.

After the low-quality cells and features were filtered, we size-normalized the gene and protein counts separately, such that each cell’s counts sum to 10,000 counts per cell. Then we log-transformed the size-normalized counts. For peaks, we binarized them by replacing all nonzero values with a value of 1. Preprocessing external single-cell expression datasets with different measurements for transfer learning and model evaluation required computing the median size factor for observed measurements from the training set and performing log-normalization afterward.

### Probabilistic modeling of multimodal datasets

Inspired by recent advancement on conditional variational inference [15, 25, 14] in the machine learning community, we aim to model the missing features and missing modalities problem altogether as a conditional probability estimation problem. Consider *m* modalities measured in the single cells. For each cell, we denote its measurement by 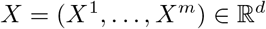 where 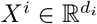 are samples of *i*th modality with *d* = *d*_1_ + … + *d_m_*. We introduce a binary mask *M* ∈ {0,1}^*d*^ for *X* and its bitwise complement *M^c^*, such that the *j*th entry of the observed sample *X_M^c^_* is *X_j_* if *M_j_* = 1 and 0 otherwise. Then, *X_M_* is defined to be *X* − *X_M^c^_*. The authentic missing pattern *M_a_* represents which components of *X* are actually missing, while the distribution of *M* can be arbitrary during training. For example, if we want to model missing completely at random, the entries of *M* could be independent Bernoulli random variable. Furthermore, we can incorporate extra structural information to model the situation of missing modality. To model the conditional distribution of the observed modalities given the missing values or modalities, we consider the following maximum likelihood problem:

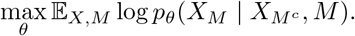

In other words, we aim to determine the conditional distribution of *X_M_* given *X_M^c^_* and *M*. Since there are totally 2^*d*^ missing patterns, we can learn 2^*d*^ conditional distributions of *X_M_*|*X_M^c^_*, *M* separately, each of them is a function: 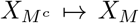 for a given *M*. However, this is computationally infeasible. Instead, we jointly model all conditional distributions by using a single neural network. Since neural networks are universal approximators [13, 14], a neural network can well approximate arbitrarily complex function: 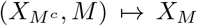 when it has enough capacity and non-polynomial activation functions. Therefore, we expect that a single neural probabilistic model can approximate *all* conditional distributions of unobserved features conditioned on any subset of observed features.

Since the above condition density itself is hard to formulate and optimize, we follow the variational Bayesian approach [4] to maximize the negative evidence lower bound (ELBO) instead:

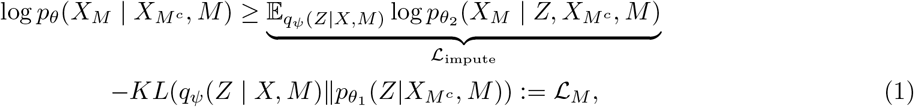

where 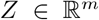 is a latent variable with approximate posterior distribution *q_ψ_*, *KL* denotes the Kullback–Leibler divergence, and *θ* = (*θ*_1_, *θ*_2_). We specify the distributions for data as follows.

In this paper, we consider trimodal analysis that includes gene expressions, protein abundances and chromatin accessibility, though the method is readily applied for more general settings. The gene counts for the nth cell are represented by a *G*-dimensional vector 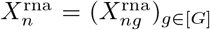 such that 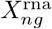 is the observed RNA count of gene *g* in cell *n*. Likewise, an A-dimensional protein counts vector 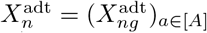 denotes the observed protein counts and an *P*-dimensional binary vector 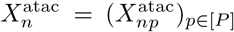 representing the occurrence of peaks for cell *n*. Let 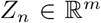 and *M_n_* be the associated joint latent variable and mask for cell *n*.

Under the target distribution *p*_*θ*_1__, we assume that the latent variables are normally distributed:

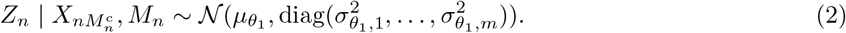

Ideally, we want *Z_n_* generated from *p*_*θ*_1__ to be as close as possible to the one generated from the the proposal distribution *q_ψ_* when *X_n_* is fully observed except for its authentic missing entries:

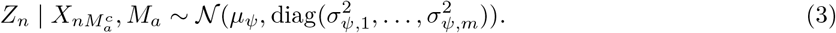

This formulation also allows us to compute the KL divergence analytically in the ELBO (1), while it is possible to extend to normal mixtures to model more complex latent structures [8]. In our implementation, we simply set 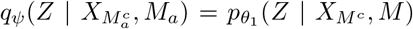 to reduce computational complexity. Finally, *q_ψ_* and *p*_*θ*_2__ are modeled as two fully-factorized Gaussian distributions, whose mean and variance are estimated by two neural networks respectively. The generative distribution *p*_*θ*_1__ are also assumed to be fully-factorized for *X_M_* given *Z*, *X_M^c^_* and *M*. We use negative binomial (NB) distribution to model the gene expression and the protein abundance. We assume that the counts are generated based on *Z_n_* as follow

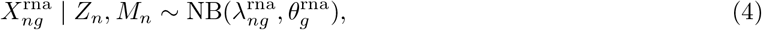

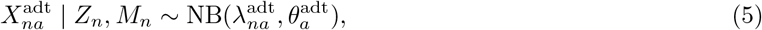

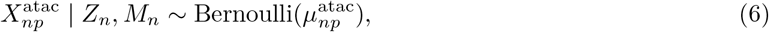

which are independent of *M* given *Z_n_*. Here the parameters 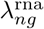 and 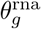 are the expected total count and the inverse dispersion of the negative binomial distribution, and the parameters 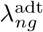 and 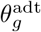 are defined analogously for protein counts. For each peak, 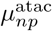 represents its posterior mean. The posterior expectations 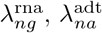, and 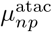 are outputted by the decoder, while the dispersion parameters are treated as trainable variables. These parameters are learned from the data.

The aforementioned probabilistic modeling (1) emphasizes missing features and modalities imputation. On the other hand, we not only want to impute the unobserved quantities, but also want to denoise the observed quantities. Therefore, we also attempt to maximize the reconstruction likelihood

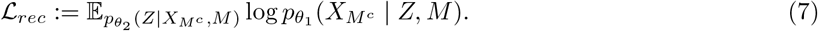

### Network architecture

scVAEIT is implemented using the Tensorflow [1] (version 2.4.1) Python library. scVAEIT consists of three main branches, the mask encoder, the main encoder and the main decoder. For each cell n, a missing mask *M_n_* is embeded as *E_n_* to a short dense vector through the mask encoder, which greatly reduces the input dimension to the main encoder and decoder. Then, the encoder takes data *X_n_* (log-normalized gene and protein counts, and binary peaks), a mask embedding vector *E_n_* and (optional) covariates *C_n_* as input, and output the estimated posterior mean and variance of the distribution of the latent variable *Z_n_*. Next, a realization is draw from this posterior distribution and fed to the decoder along with the mask embedding vector *E_n_* and the covariates *C_n_*. The decoder finally outputs the posterior mean of *X*.

We use subconnections at the first layer of the encoder and the last layer of the decoder. In each of these layer, there are 256 and 128 units for genes and proteins respectively, and 16 units for peaks in each chromosome. The weights are isolated between different blocks. The mask encoder outputs 32-dim, 16-dim, and 2-dim vectors for genes, proteins, and peaks in each chromosome. Besides these special layers, we also have one fully-connected hidden layer of 256 units in the encoder and the decoder, with LeakyReLU activation functions.

### Model training

scVAEIT is trained in an end-to-end manner. The objective function is a convex combination of the ELBO (1) and the reconstruction likelihood (7):

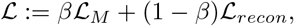

where *β* ∈ [0,1] is a hyperparameter set to be 0.5 for all experiments. That is, the parameters are optimized by Monte Carlo sampling to maximize the weighted average of the reconstruction likelihood and the imputation likelihood, while minimizing the KL divergence between masked posterior latent variable *Z*|*X_M^c^_*, *M* and the authentic posterior latent variable 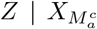, *M_a_*. During training, with equal probability we observe the original data and the masked data. The mask is repeatedly randomly generated for each cell at the beginning of every gradient update step in each epoch during the optimization process, such that each modality is observed with equal probability, and each entry is further randomly masked out with probability 0.2. The sensitivity analysis on the masking probability is also included in Fig.S8. To balance the magnitudes of different modalities, we calculated the weighted likelihood using weights 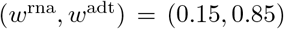 for bimodal datasets and 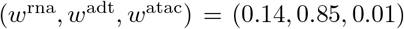. The default variable initializer in Tensorflow is used, sampling weight matrix from a uniform distribution and setting bias vectors to be zero. We train our model for 300 epochs using the AdamW optimizer [19], a variant of the stochastic gradient descent algorithm, with a batch size of 512, a learning rate of 10^−3^, and a weight decay of 10^−4^. We also use batch normalization to aid in training stability. Because we use mini-batches for training, scVAEIT’s memory usage does not effected by the number of cells. Instead, it is only related to in the number of features in the dataset and number of neural network parameters.

### Benchmarking methods

We compare scVAEIT with Seurat[12], totalVI[9], and MultiVI[3]. Seurat v4’s weighted nearest neighbor (WNN) is used to perform intermediate integration and multimodal anchor-based transfer-learning method is used to perform transfer learning to new datasets. Standard preprocessing procedures in Seurat are used for evaluating Seurat’s results. More specifically, RNA counts are normalized by LogNormalization method and protein counts are normalized by centered log-ratio (CLR) method. When evaluating Seurat’s protein imputation result, we revert the CLR normalized imputed counts and perform log-normalization. The log-normalized gene counts and size-normalized protein counts are provided as input to totalVI; the log-normalized gene counts and binary peaks are provided as input to MultiVI. For stimulation effect correction, we use Harmony[16]‘s corrected dimension reductions for running Seurat’s WNN, and provide an indicator variable as a covariate to totalVI and MultiVI. For running Harmony and 3-WNN integration on DOGMA-seq datasets, we use the script provided in the original paper[23] (https://github.com/caleblareau/asap_reproducibility/blob/master/pbmc_stim_multiome/code/11_setup.R). For running UINMF[17] (version 1.1.0) on the three multimodal datasets, we first split each dataset into two based on stimulation conditions. Then each of the six datasets are preprocessed with functions `normalize` and `scaleNotCenter` according to their tutorials. After that, the shared features and unshared features of the six datasets are separated and supplied to function optimizeALS, where the parameters are chosen by examining the results of functions `suggestK` and `suggestLambda`. The imputed values are obtained with function `imputeKNN`. For running totalVI and MultiVI, we set the latent dimension as 32, `early_stopping_patience` as 15 and leaving all other hyperparameters as default. All code required to reproduce our reported results, including data preprocessing and model training, have been deposited on GitHub (https://github.com/jaydu1/scVAEIT).

### Evaluation metric

We use multiple evaluation metrics for comparing different methods on imputing gene counts, protein counts and peaks. As the log-normalized gene and protein expressions are treated as continuous, we use Pearson correlation and Spearman correlation to evaluate their imputation quality. For *n* observations (*x*_1_,…, *x_n_*) and their prediction/imputation values (*y*_1_,…, *y_n_*), the correlation metrics are defined as

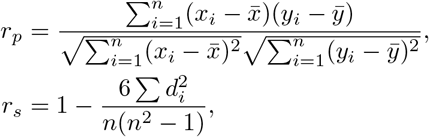

where 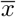 and 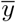 are the mean of *x_i_*’s and *y_i_*’s respectively, and *d_i_* = rank(*x_i_*) − rank(*y_i_*) is the difference between the two ranks of each observation. The imputation error is also quantified by the root mean square error (RMSE),

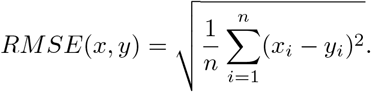

For chromatin accessibility, the peaks are binarized while the imputed values are continuous in [0,1]. Thus we use area under the receiver operating characteristic (AUROC), binary cross entropy (BCE) and RMSE to evaluate imputation accuracy of binary peaks. The AUROC metric takes value between 0 and 1, which is commonly used in statistics and machine learning community. A larger value of AUROC indicates that the binary outcome is easier to predict based on imputed value at various threshold settings. The BCE metric

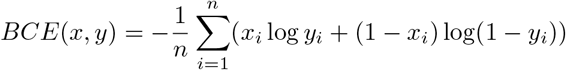

is equivalent to the negative log-likelihood of Bernoulli variables. Thus a smaller value of BCE means a better fit of statistical models. For each evaluation metric, we effectively consider each gene, protein, or peak in each cell a separate observation.

## Data availability

All datasets used in this paper are previously published and freely available. For bimodal datasets integration, CITE-seq PBMC cells from Hao et al.[12] are in the GEO database under accession code GSE164378 and a Seurat object containing filtered cells is also provided in their tutorial https://satijalab.org/seurat/articles/multimodal_reference_mapping.html; the CITE-seq CBMC cells from Stoeckius et al.[26] are in the GEO database under accession code GSE100866 and a Seurat object containing these cells is also available as ‘bmcite’ in the SeuratData (v0.2.1) package; the REAP-seq PBMC cells from Peterson et al.[24] are in the GEO database under accession code GSE100501. For trimodel datasets integration, the DOGMA-seq, CITE-seq, and ASAP-seq PBMC cells from Mimitou et al.[23] are in the GEO database under accession code GSE156478, and the intermediate result files are retrieved from their Github repository https://github.com/caleblareau/asap_reproducibility.

## Code availability

The Python package of scVAEIT is publicly available at https://github.com/jaydu1/scVAEIT with MIT license. Python and R scripts for reproducing all results in this paper are also provided in the same repository.

## Acknowledgements

This work used the Bridges-2 system, which is supported by NSF award number OAC-1928147 at the Pittsburgh Supercomputing Center (PSC)[30, 6] This project was funded by National Institute of Mental Health (NIMH) grant R01MH123184.

## Author contributions statement

J.D., Z.C., and K.R. conceived the experiment(s). J.D. conducted the experiment(s). J.D., Z.C., and K.R. analysed the results. J.D., Z.C., and K.R. reviewed the manuscript.

## Supplement

### Datasets

**Table S1:** Summary table of datasets analyzed in this study, with only highly variable features included. The numbers of the shared features to the CITE-seq PBMC[12] dataset are given in parentheses.

| Dataset | Number of Measurements |  |  |  |
| --- | --- | --- | --- | --- |
|  | Cell | Gene | Protein | Peak |
| CITE-seq PBMC[12] | 161764 | 4686 | 227 |  |
| CITE-seq CBMC[26] | 7891 | 14348(3464) | 10(10) |  |
| REAP-seq PBMC[24] | 7092 | 32738(3864) | 43(38) |  |
| DOGMA-seq PBMC[23] | 13763 | 2166 | 208 | 26765 |
| ASAP-seq PBMC[23] | 8535 |  | 227 | 38152 |
| CITE-seq PBMC[23] | 8689 | 2374 | 227 |  |
| Total | 30987 | 3660 | 227 | 38711 |

### The evidence lower bound

The derivation for variational lower bound (1) of the conditional log-likelihood is given by

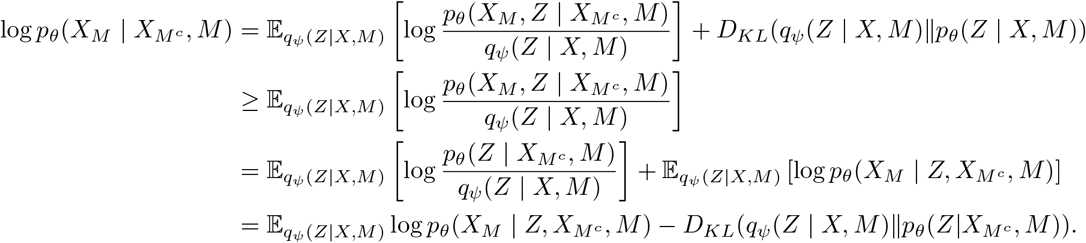

### The network architecture

**Table S2:** Comparison of the performance of scVAEIT on the held-out cell type Mono with varying latent dimension sizes. The 32-dimensional latent space has slightly better performance compared to the 16-dimensional latent space, while it is similar to the most complex setting when we use a 64-dimensional latent space. Hence, we elect to train scVAEIT using a 32-dimensional latent space.

| Metric | Source | Target | Dimension of Latent Space |  |  |
| --- | --- | --- | --- | --- | --- |
|  |  |  | 16 | 32 | 64 |
| Pearson's $r$ | ADT | ADT | 0.90 | 0.91 | 0.91 |
|  |  | RNA | 0.80 | 0.81 | 0.80 |
|  | RNA | ADT | 0.88 | 0.87 | 0.89 |
|  |  | RNA | 0.82 | 0.83 | 0.83 |
| Spearman's $r$ | ADT | ADT | 0.89 | 0.90 | 0.89 |
|  |  | RNA | 0.54 | 0.55 | 0.54 |
|  | RNA | ADT | 0.85 | 0.84 | 0.86 |
|  |  | RNA | 0.56 | 0.56 | 0.56 |

### Supplementary results

**Figure S1:**
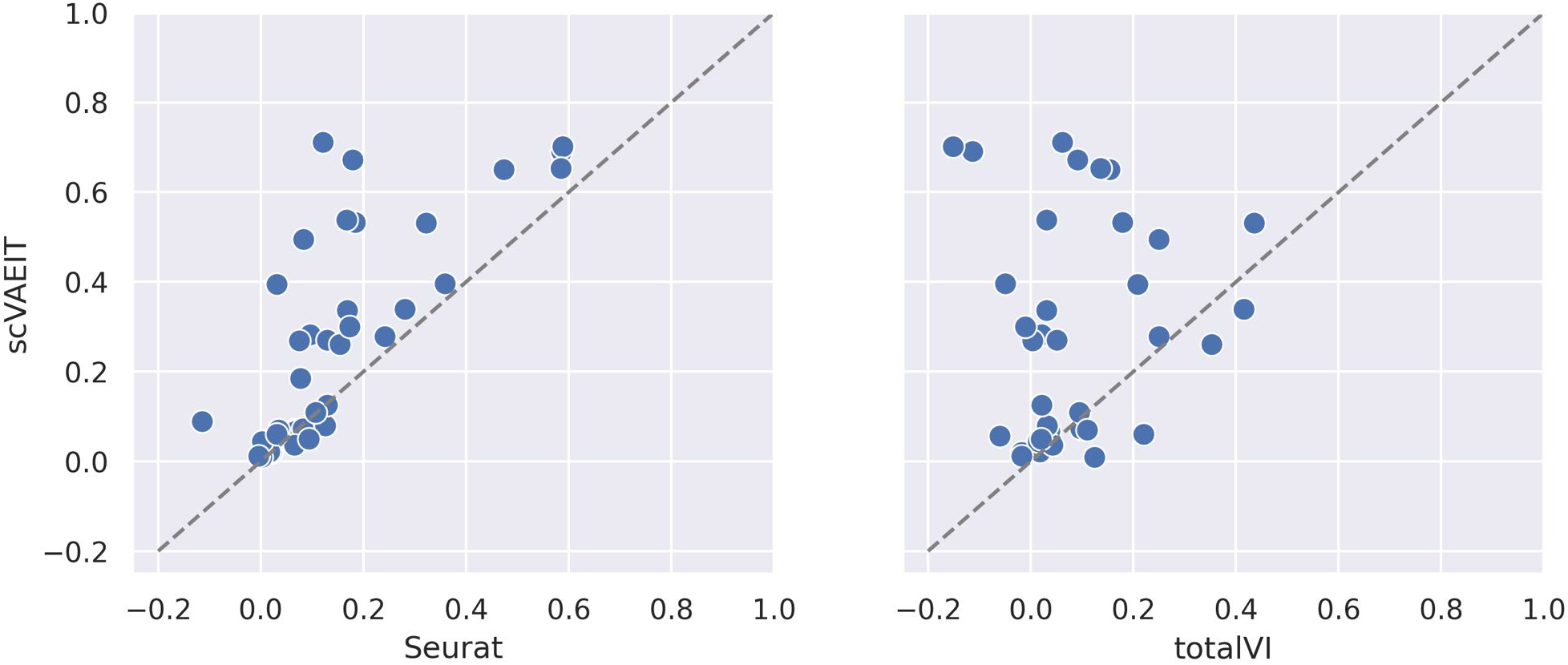
Across technique correlations between imputed and measured protein abundance on the REAP-seq PBMC data: (a) scVAEIT versus Seurat and (b) scVAEIT versus totalVI.

**Figure S2:**
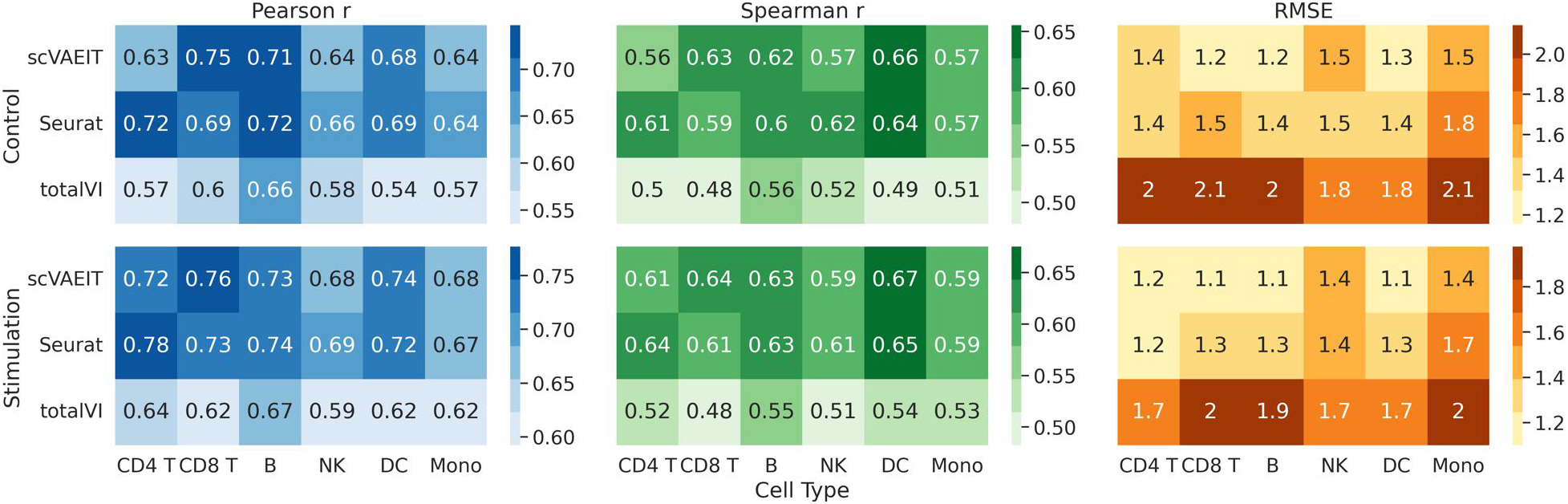
Performance of protein imputation on the held-out cell types of the DOGMA-seq dataset.

**Figure S3:**
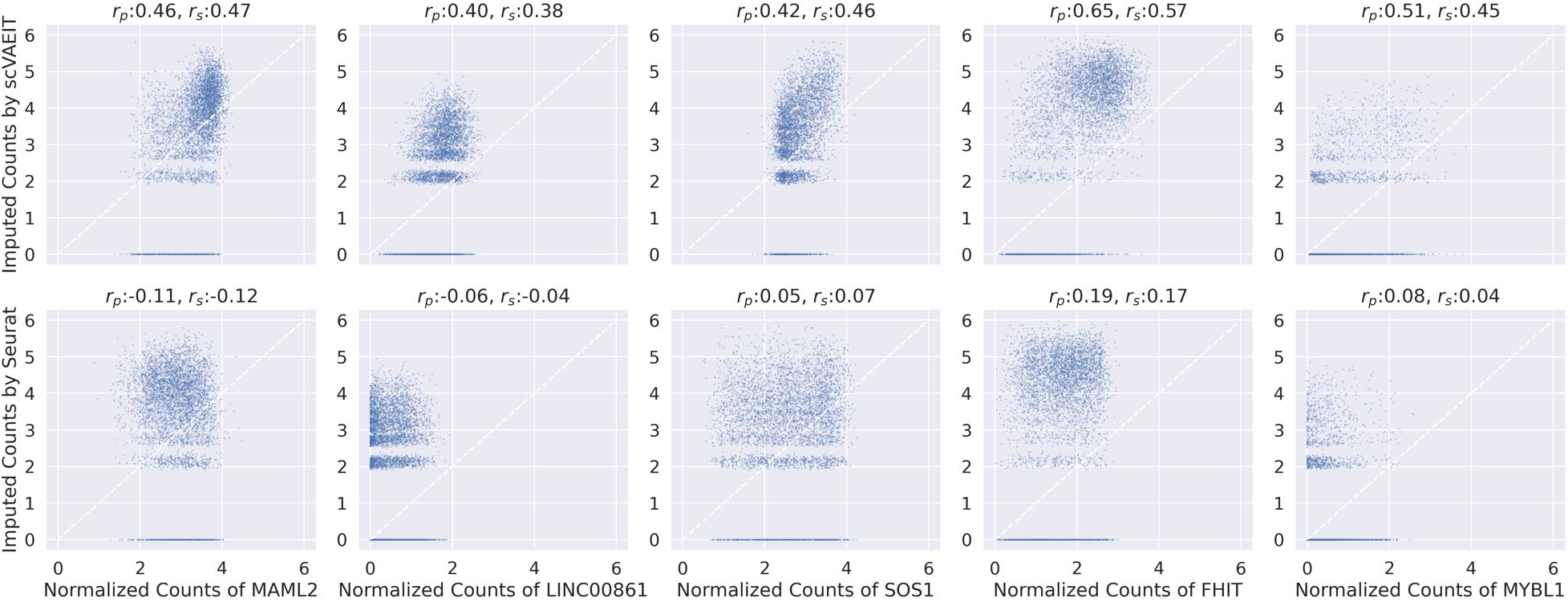
Scatter plot of normalized gene counts versus imputed values on the held-out CD4 T cell type of the DOGMA-seq dataset by scVAEIT and Seurat.

**Figure S4:**
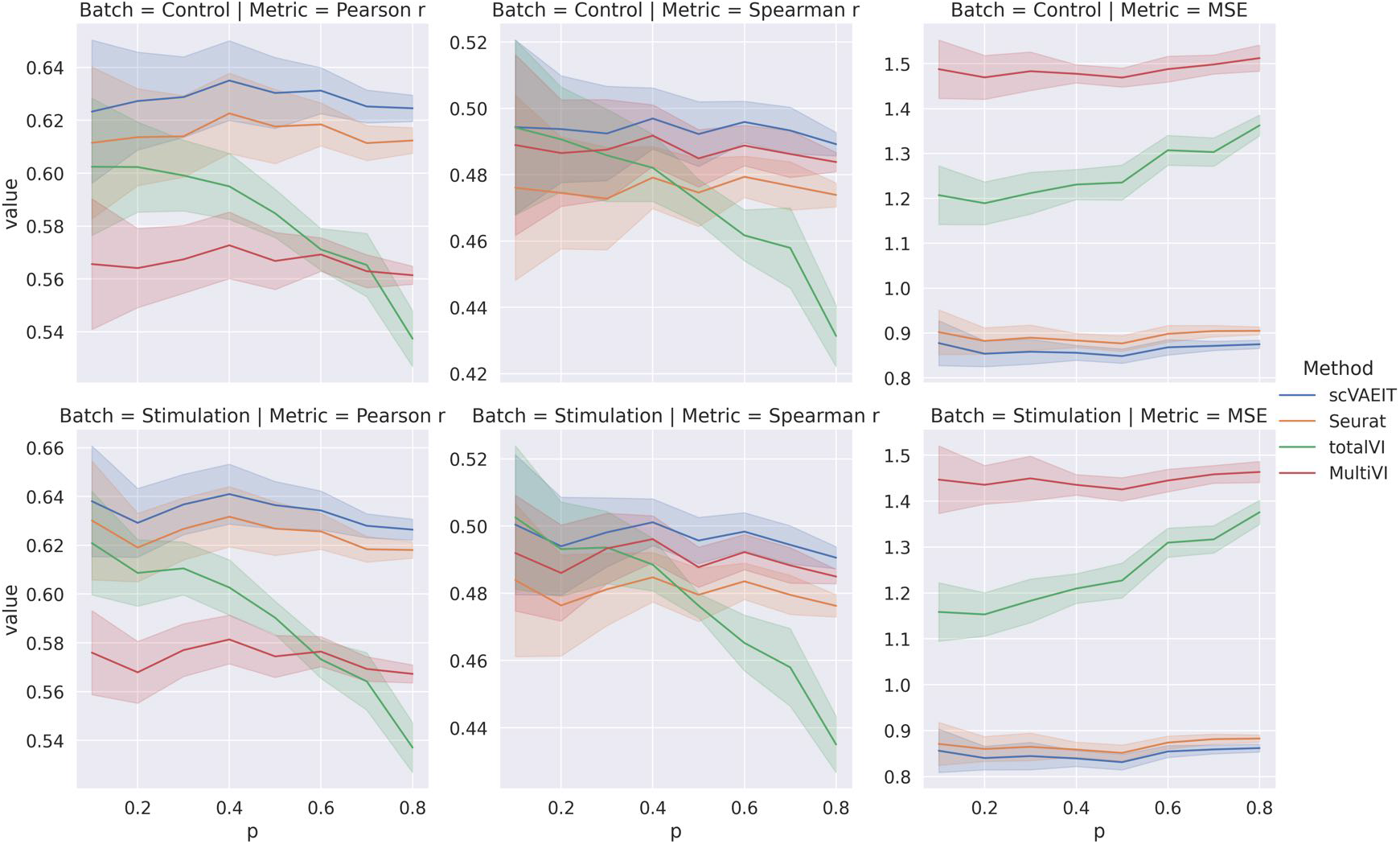
Performance of gene imputation on the held-out CD4 T cell type of the DOGMA-seq dataset with random missing. For each missing proportion, all methods are evaluated using the same observed features across 10 runs and the shaded region is within one standard deviation of the mean.

**Figure S5:**
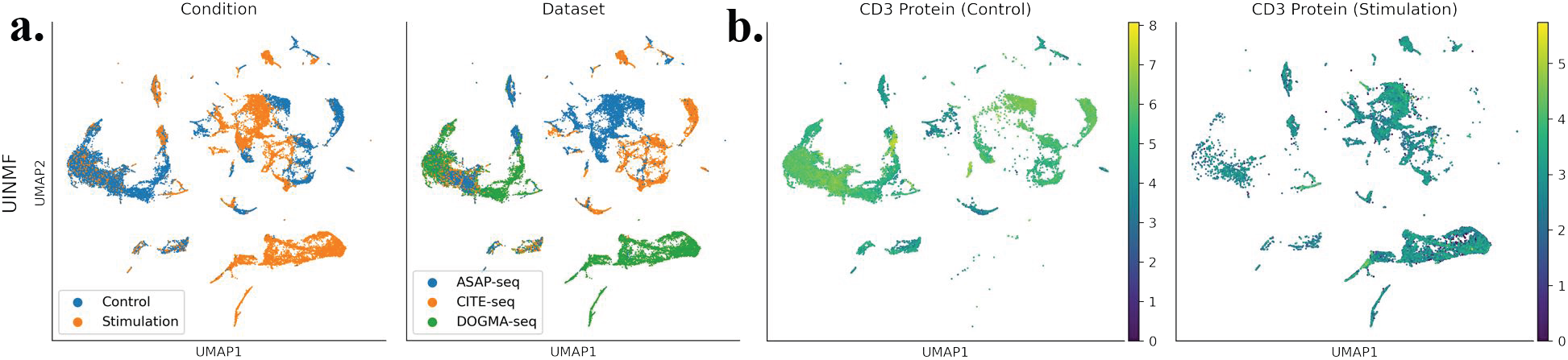
Integration of DOGMA-seq, CITE-seq, and ASAP-seq PBMC datasets by UINMF. **(a)** Joint embeddings of the three multimodal datasets after intermediate integration. **(b)** Log-normalized expressions of CD3 protein in the control cells (first column) and the stimulated cells (second column) on scVAEIT’s and Seurat’s embeddings. Purple and yellow dots correspond to lowly and highly expressed cells, respectively.

**Figure S6:**
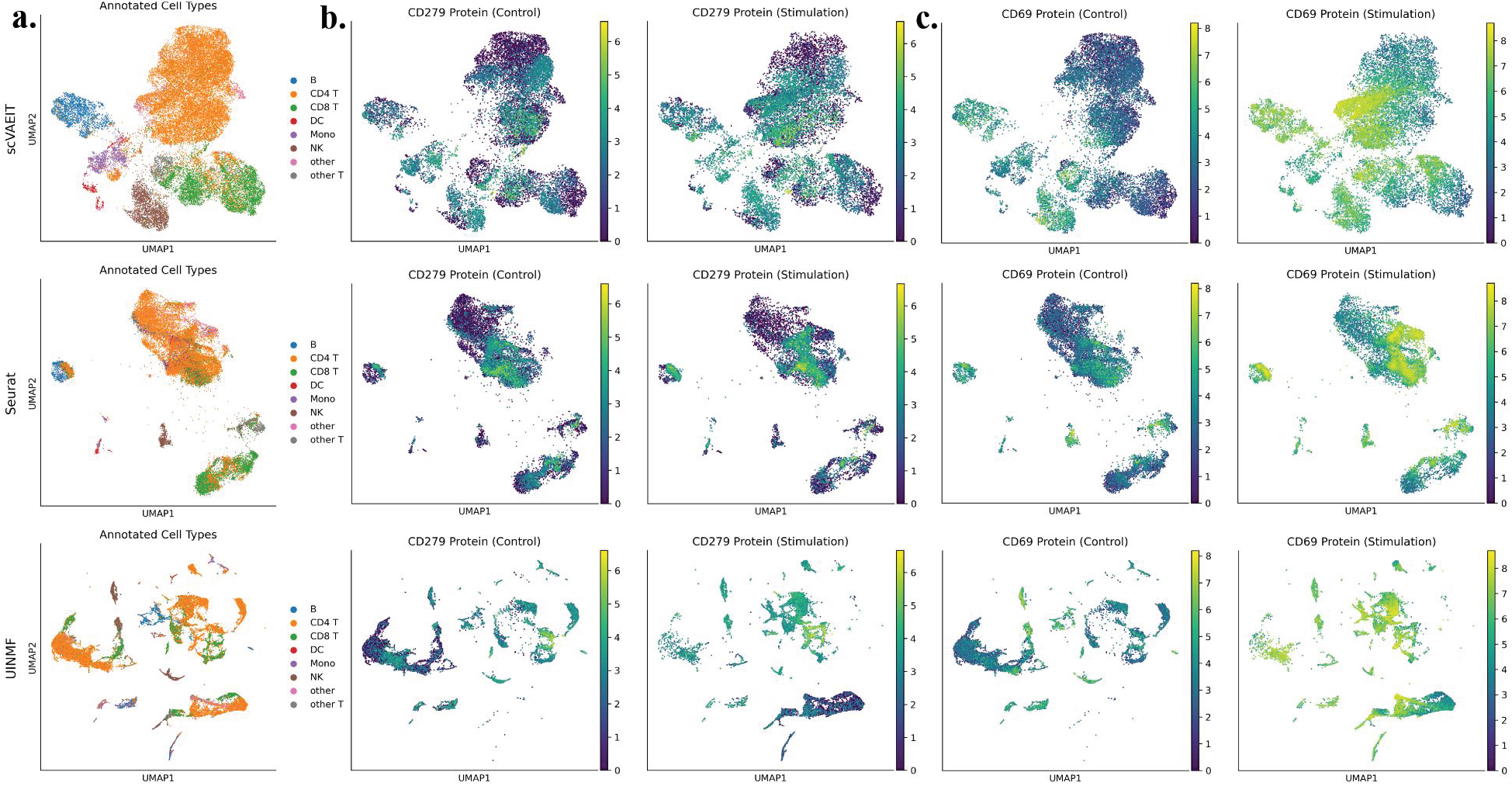
Integration of DOGMA-seq, CITE-seq, and ASAP-seq PBMC datasets. **(a)** Joint embeddings of the three multimodal datasets after intermediate integration, colored by the annotated cell types. **(b-c)** Log-normalized expressions of **(b)** CD279 protein and **(c)** CD69 protein in the control cells (first column) and the stimulated cells (second column) on scVAEIT’s, Seurat’s, and UINMF’s embeddings. Purple and yellow dots correspond to lowly and highly expressed cells respectively.

**Figure S7:**
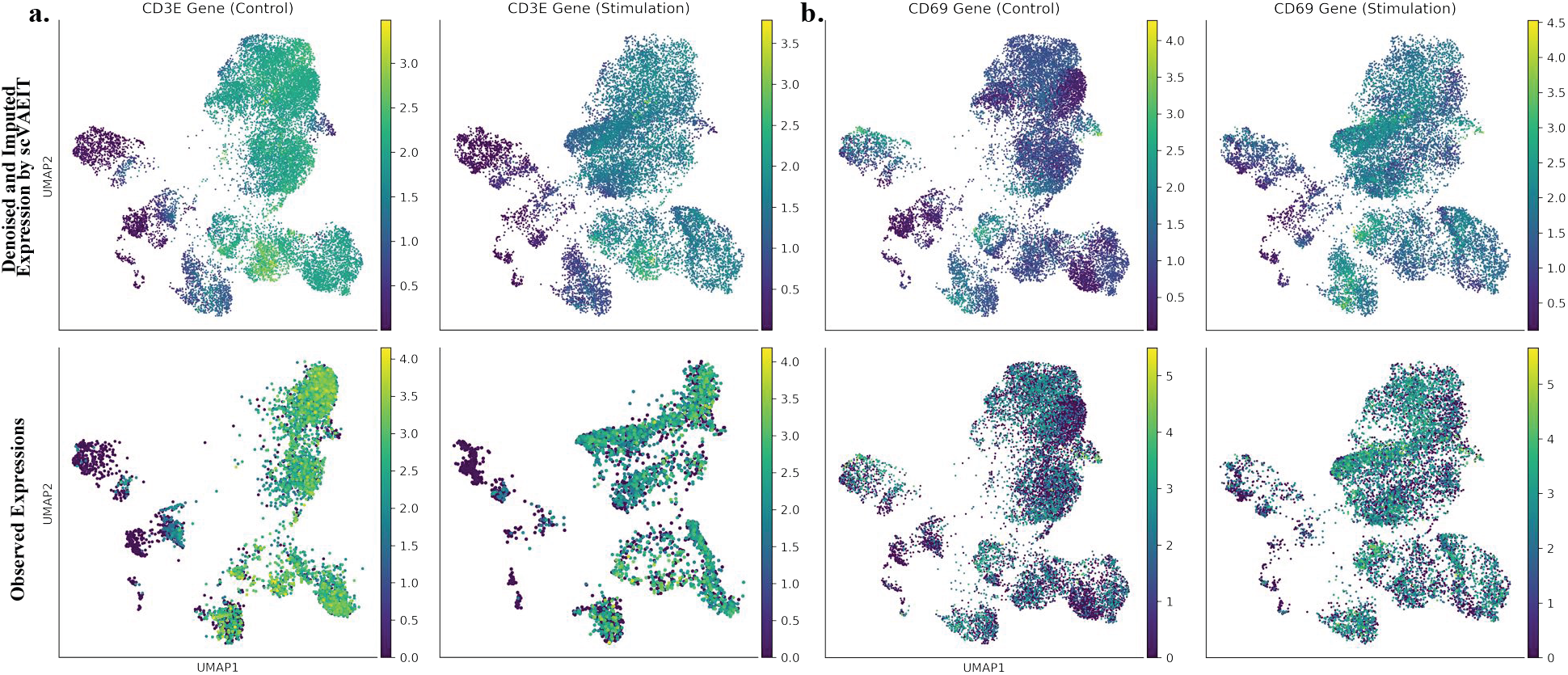
Gene expressions after integration of DOGMA-seq, CITE-seq, and ASAP-seq PBMC datasets by scVAEIT. **(a)-(b)** Joint embeddings of the three multimodal datasets after intermediate integration, colored by the log-normalized expressions of **(a)** CD3E gene and **(b)** CD69 gene in the control cells (first column) and the stimulated cells (second column). The first row shows the denoised and imputed expressions by scVAEIT, and the second row shows the observed expressions. The CD3E gene is only observed in the CITE-seq dataset, while the CD69 gene is only observed in the DOGMA-seq and CITE-seq datasets. Purple and yellow dots correspond to lowly and highly expressed cells respectively.

**Figure S8:**
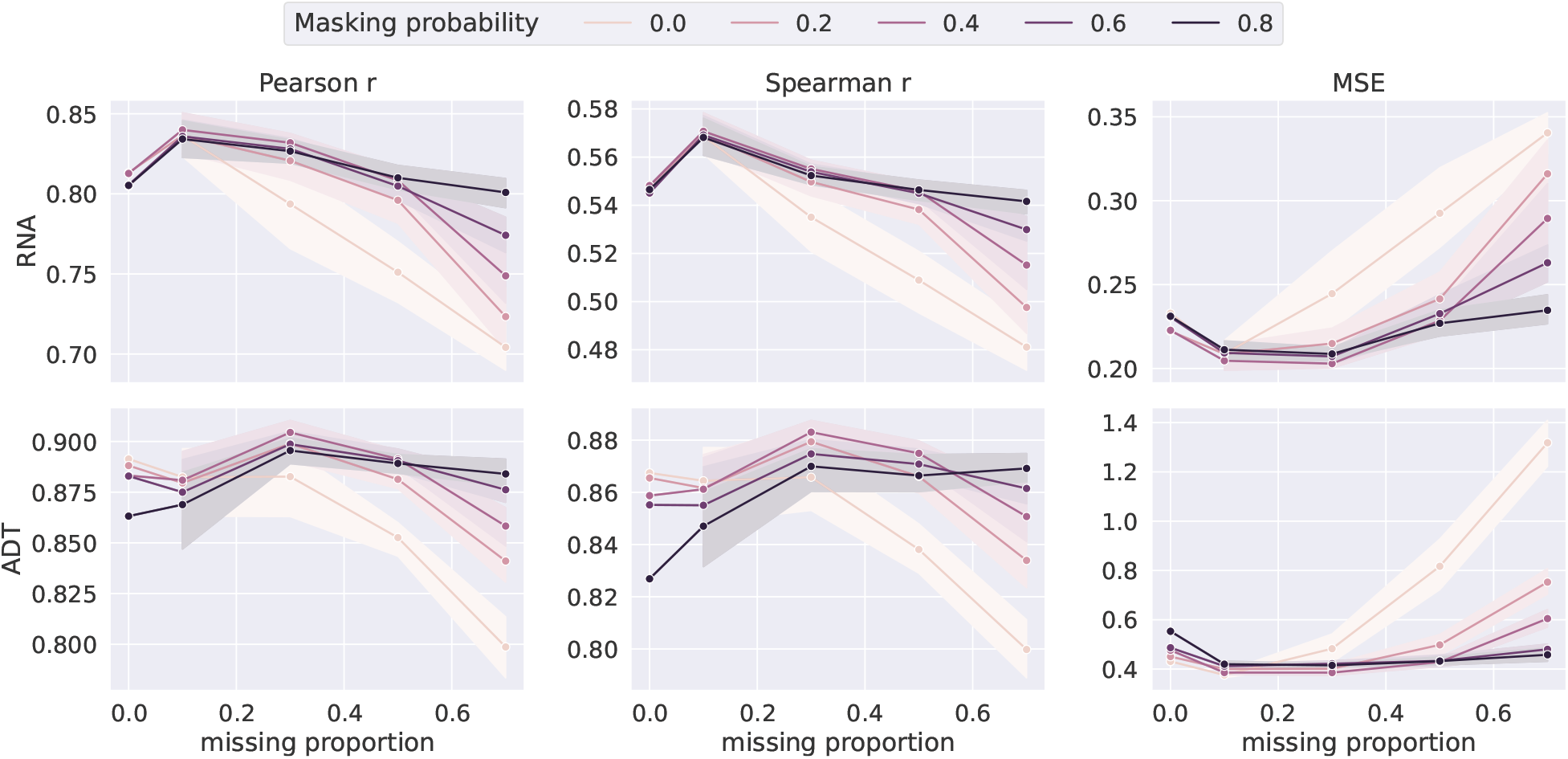
Sensitivity analysis of the masking probability for scVAEIT. With masking probability varying from 0 to 0.8, scVAEIT is trained on the PBMC CITE-seq dataset [12] with the Mono cell type held out for evaluation. The three columns correspond to the Pearson *r*, the Spearman *r* and the MSE (mean square error) for cross-modal translation from proteins to genes (the first row) and from genes to proteins (the second row), with random missing. ADT stands for antibody-derived tags for measuring proteins. For each missing proportion, the method is evaluated across 10 runs and the shaded region is within one standard deviation of the average performance. When the missing proportion is small, scVAEIT behaves similarly for various masking probabilities; when the missing proportion increases, scVAEIT trained with higher masking probability has better performance.

**Figure S9:**
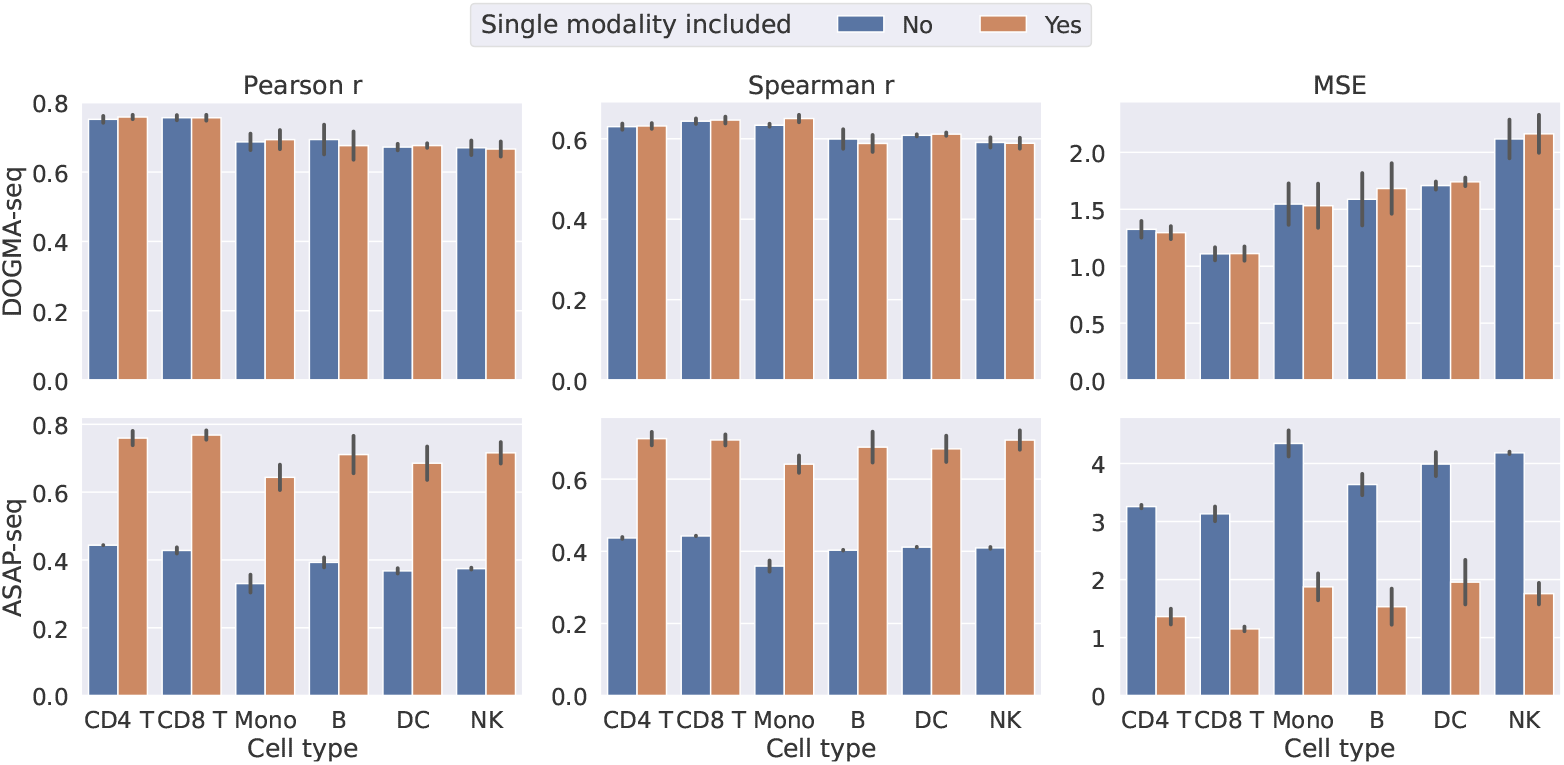
Performance of protein imputation on the held-out cell types, when integrating multimodal datasets with and without single-modal datasets. The Pearson *r*, Spearman *r*, and MSE (mean square error) on the control and stimulated groups are visualized. The baseline scVAEIT models without single modality included are trained on bimodal datasets containing the genes and proteins from the PBMC DOGMA-seq dataset [23] with one cell type excluded at a time; the compared models are trained on both the above bimodal datasets and the corresponding single-modal datasets containing the proteins from the PBMC ASAP-seq dataset [23]. The inclusion of single-modal datasets did not hurt the performance on imputing the bimodal datasets, and even slightly improves the performance. On the other hand, there is substantial benefit when imputing the single-modal dataset and the improved performance is similar to that of bimodal datasets.

**Figure S10:**
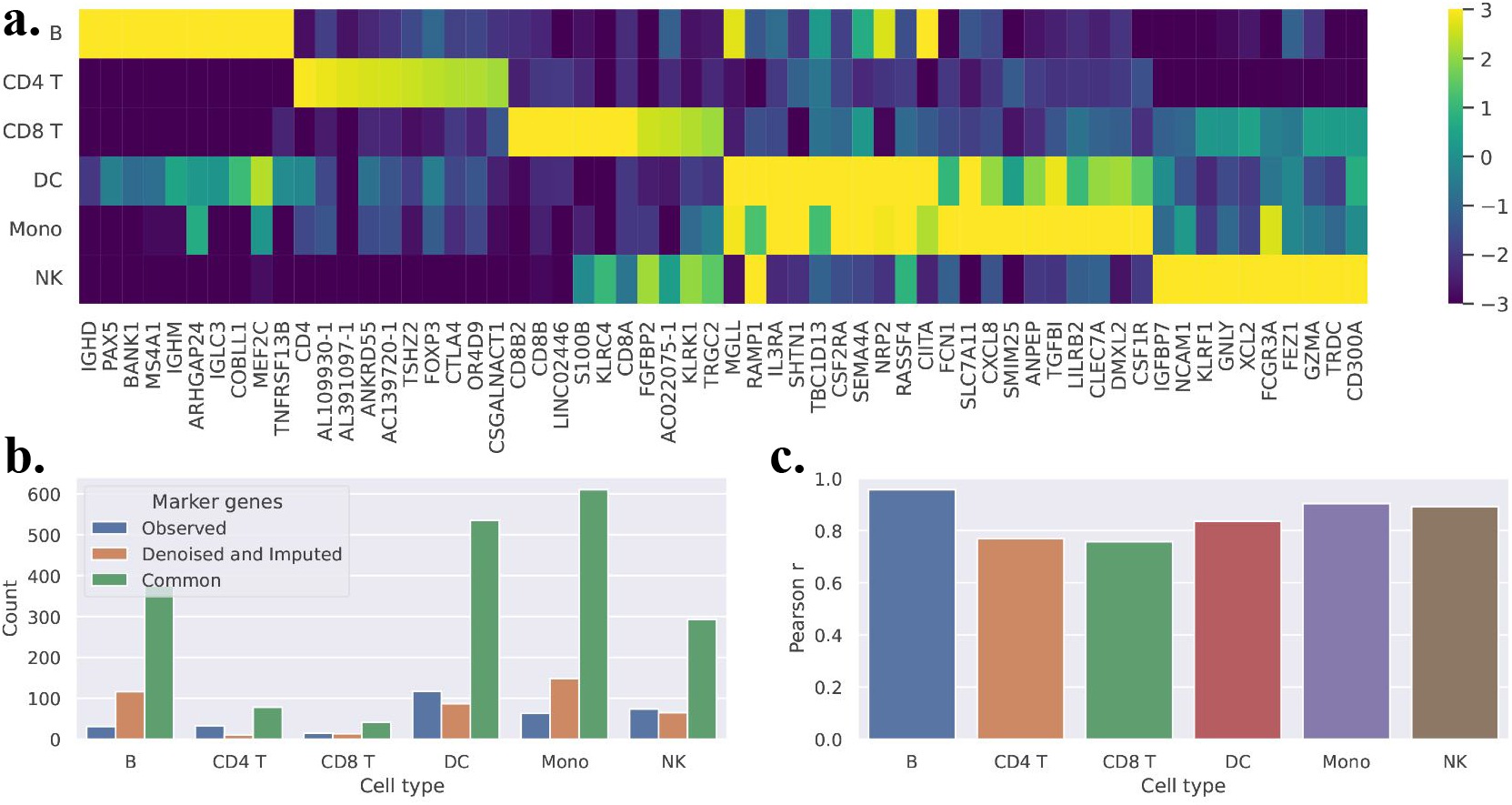
Differential expressed testing with scVAEIT on the PBMC DOGMA-seq, CITE-seq, and ASAP-seq datasets [23]. **(a)** Heatmap of log_2_(fold changes) of top 10 upregulated denoised and imputed genes in each cell type (Wilcoxon Rank Sum test, adjusted *p*-value< 0.001 with Bonferroni correction). The fold change of a gene in a cluster is defined to be the ratio of the average expression in this cluster to the one outside of the cluster. **(b)** Barplots of counts of marker genes identified based on observed expression, and denoised and imputed expression (*p*-value< 0.001 and log_2_(fold changes)> 1). The exclusive counts and the common/intersected counts are visualized. The common marker genes compose a high proportion of both the two sets of marker genes, meaning that most marker genes of the observed data can be discovered by testing on the imputed data. **(c)** The Pearson *r* of the observed and denoised log_2_(fold changes) on all genes. There are high correlations between the two among all cell types, especially for non-T cell types.

## References

[1] M. Abadi, A. Agarwal, P. Barham, E. Brevdo, Z. Chen, C. Citro, G. S. Corrado, A. Davis, J. Dean, M. Devin, S. Ghemawat, I. Goodfellow, A. Harp, G. Irving, M. Isard, Y. Jia, R. Jozefowicz, L. Kaiser, M. Kudlur, J. Levenberg, D. Mané, R. Monga, S. Moore, D. Murray, C. Olah, M. Schuster, J. Shlens, B. Steiner, I. Sutskever, K. Talwar, P. Tucker, V. Vanhoucke, V. Vasudevan, F. Viégas, O. Vinyals, P. Warden, M. Wattenberg, M. Wicke, Y. Yu, and X. Zheng. TensorFlow: Large-scale machine learning on heterogeneous systems, 2015. URL https://www.tensorflow.org/. Software available from tensorflow.org.

[2] R. Argelaguet, A. S. Cuomo, O. Stegle, and J. C. Marioni. Computational principles and challenges in single-cell data integration. Nature biotechnology, 39(10):1202–1215, 2021.

[3] T. Ashuach, M. I. Gabitto, M. I. Jordan, and N. Yosef. Multivi: deep generative model for the integration of multi-modal data. bioRxiv, 2021.

[4] D. M. Blei, A. Kucukelbir, and J. D. McAuliffe. Variational inference: A review for statisticians. Journal of the American statistical Association, 112(518):859–877, 2017.

[5] P. Boyeau, J. Regier, A. Gayoso, M. I. Jordan, R. Lopez, and N. Yosef. An empirical bayes method for differential expression analysis of single cells with deep generative models. bioRxiv, 2022.

[6] S. T. Brown, P. Buitrago, E. Hanna, S. Sanielevici, R. Scibek, and N. A. Nystrom. Bridges-2: a platform for rapidly-evolving and data intensive research. In Practice and Experience in Advanced Research Computing, pages 1–4. Association for Computing Machinery, 2021.

[7] S.-K. Chu, S. Zhao, Y. Shyr, and Q. Liu. Comprehensive evaluation of noise reduction methods for single-cell RNA sequencing data. Briefings in Bioinformatics, 23(2), jan 2022.

[8] J.-H. Du, M. Gao, and J. Wang. Model-based trajectory inference for single-cell rna sequencing using deep learning with a mixture prior. bioRxiv, 2020.

[9] A. Gayoso, Z. Steier, R. Lopez, J. Regier, K. L. Nazor, A. Streets, and N. Yosef. Joint probabilistic modeling of single-cell multi-omic data with totalvi. Nature methods, 18(3):272–282, 2021.

[10] S. Ghazanfar, C. Guibentif, and J. C. Marioni. Stabmap: Mosaic single cell data integration using non-overlapping features. bioRxiv, 2022.

[11] L. Haghverdi, A. T. Lun, M. D. Morgan, and J. C. Marioni. Batch effects in single-cell rna-sequencing data are corrected by matching mutual nearest neighbors. Nature biotechnology, 36(5):421–427, 2018.

[12] Y. Hao, S. Hao, E. Andersen-Nissen, W. M. Mauck III, S. Zheng, A. Butler, M. J. Lee, A. J. Wilk, C. Darby, and M. Zager. Integrated analysis of multimodal single-cell data. Cell, 184(13):3573–3587, 2021.

[13] K. Hornik, M. Stinchcombe, and H. White. Multilayer feedforward networks are universal approximators. Neural networks, 2(5):359–366, 1989.

[14] O. Ivanov, M. Figurnov, and D. Vetrov. Variational autoencoder with arbitrary conditioning. In International Conference on Learning Representations, 2018.

[15] D. P. Kingma and M. Welling. Auto-encoding variational bayes. In Y. Bengio and Y. LeCun, editors, 2nd International Conference on Learning Representations, 2014. URL http://arxiv.org/abs/1312.6114.

[16] I. Korsunsky, N. Millard, J. Fan, K. Slowikowski, F. Zhang, K. Wei, Y. Baglaenko, M. Brenner, P.-r. Loh, and S. Raychaudhuri. Fast, sensitive and accurate integration of single-cell data with harmony. Nature methods, 16(12):1289–1296, 2019.

[17] A. R. Kriebel and J. D. Welch. Uinmf performs mosaic integration of single-cell multi-omic datasets using nonnegative matrix factorization. Nature communications, 13(1):1–17, 2022.

[18] R. Lopez, P. Boyeau, N. Yosef, M. Jordan, and J. Regier. Decision-making with auto-encoding variational bayes. Advances in Neural Information Processing Systems, 33:5081–5092, 2020.

[19] I. Loshchilov and F. Hutter. Decoupled weight decay regularization. In International Conference on Learning Representations, 2017.

[20] M. D. Luecken, M. Büttner, K. Chaichoompu, A. Danese, M. Interlandi, M. F. Müller, D. C. Strobl, L. Zappia, M. Dugas, and M. Colomé-Tatché. Benchmarking atlas-level data integration in single-cell genomics. Nature methods, 19(1):41–50, 2022.

[21] L. McInnes, J. Healy, N. Saul, and L. Großberger. Umap: Uniform manifold approximation and projection. Journal of Open Source Software, 3(29):861, 2018.

[22] Z. Miao, B. D. Humphreys, A. P. McMahon, and J. Kim. Multi-omics integration in the age of million single-cell data. Nature Reviews Nephrology, 17(11):710–724, 2021.

[23] E. P. Mimitou, C. A. Lareau, K. Y. Chen, A. L. Zorzetto-Fernandes, Y. Hao, Y. Takeshima, W. Luo, T.-S. Huang, B. Z. Yeung, E. Papalexi, P. I. Thakore, T. Kibayashi, J. B. Wing, M. Hata, R. Satija, K. L. Nazor, S. Sakaguchi, L. S. Ludwig, V. G. Sankaran, A. Regev, and P. Smibert. Scalable, multimodal profiling of chromatin accessibility, gene expression and protein levels in single cells. Nature Biotechnology, 39(10):1246–1258, jun 2021. URL https://doi.org/10.1038%2Fs41587-021-00927-2.

[24] V. M. Peterson, K. X. Zhang, N. Kumar, J. Wong, L. Li, D. C. Wilson, R. Moore, T. K. McClanahan, S. Sadekova, and J. A. Klappenbach. Multiplexed quantification of proteins and transcripts in single cells. Nature biotechnology, 35(10):936–939, 2017.

[25] K. Sohn, H. Lee, and X. Yan. Learning structured output representation using deep conditional generative models. Advances in neural information processing systems, 28, 2015.

[26] M. Stoeckius, C. Hafemeister, W. Stephenson, B. Houck-Loomis, P. K. Chattopadhyay, H. Swerdlow, R. Satija, and P. Smibert. Simultaneous epitope and transcriptome measurement in single cells. Nature methods, 14(9):865–868, 2017.

[27] T. Stuart and R. Satija. Integrative single-cell analysis. Nature reviews genetics, 20(5):257–272, 2019.

[28] T. Stuart, A. Butler, P. Hoffman, C. Hafemeister, E. Papalexi, W. M. Mauck III, Y. Hao, M. Stoeckius, P. Smibert, and R. Satija. Comprehensive integration of single-cell data. Cell, 177(7):1888–1902, 2019.

[29] E. Swanson, C. Lord, J. Reading, A. T. Heubeck, P. C. Genge, Z. Thomson, M. D. Weiss, X. jun Li, A. K. Savage, R. R. Green, T. R. Torgerson, T. F. Bumol, L. T. Graybuck, and P. J. Skene. Simultaneous trimodal single-cell measurement of transcripts, epitopes, and chromatin accessibility using TEA-seq. eLife, 10, apr 2021. URL https://doi.org/10.7554%2Felife.63632.

[30] J. Towns, T. Cockerill, M. Dahan, I. Foster, K. Gaither, A. Grimshaw, V. Hazlewood, S. Lathrop, D. Lifka, and G. D. Peterson. Xsede: accelerating scientific discovery. Computing in science & engineering, 16(5):62–74, 2014.

[31] H. T. N. Tran, K. S. Ang, M. Chevrier, X. Zhang, N. Y. S. Lee, M. Goh, and J. Chen. A benchmark of batch-effect correction methods for single-cell rna sequencing data. Genome biology, 21(1):1–32, 2020.

[32] J. Wang, D. Agarwal, M. Huang, G. Hu, Z. Zhou, C. Ye, and N. R. Zhang. Data denoising with transfer learning in single-cell transcriptomics. Nature methods, 16(9):875–878, 2019.

[33] K. E. Wu, K. E. Yost, H. Y. Chang, and J. Zou. Babel enables cross-modality translation between multiomic profiles at single-cell resolution. Proceedings of the National Academy of Sciences, 118(15), 2021.

[34] Z. Zhou, C. Ye, J. Wang, and N. R. Zhang. Surface protein imputation from single cell transcriptomes by deep neural networks. Nature Communications, 11, 12 2020. ISSN 20411723.

